# Orientation dependent proton transverse relaxation in the human brain white matter: The magic angle effect on a cylindrical helix

**DOI:** 10.1101/2021.09.13.460097

**Authors:** Yuxi Pang

**Affiliations:** Department of Radiology, University of Michigan, Ann Arbor, MI, USA

**Keywords:** cylindrical helix model, diffusion tensor imaging, quantitative magnetization transfer, magic angle effect, orientation-dependent transverse relaxation, principal diffusivity direction, white matter

## Abstract

**Purpose:** To overcome some limitations of prior orientation-dependent proton transverse relaxation formalisms in white matter (WM) with a novel framework based on the generalized magic angle effect function.

**Methods:** A cylindrical helix model was developed embracing anisotropic rotational and translational diffusion of restricted molecules in human brain WM, with the former characterized by an axially symmetric system. Transverse relaxation rates *R*_2_ and 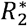 were divided into isotropic 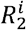 and anisotropic parts, 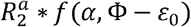, with *α* denoting an open angle and *ε*_0_ an orientation (Φ) offset from DTI-derived primary diffusivity direction. The proposed framework (Fit A) was compared with prior models without *ε*_0_ on previously published water and methylene proton transverse relaxation rates from developing, healthy, and pathological WM at 3T. Goodness of fit was represented by root-mean-square error (RMSE). *F*-test and linear correlation were used with statistical significance set to *P* ≤ 0.05.

**Results:** Fit A significantly (*P*<0.01) outperformed prior models as demonstrated by reduced RMSEs, e.g., 0.349 vs. 0.724 in myelin water. Fitted *ε*_0_ was in good agreement with calculated *ε*_0_ from directional diffusivities. Compared with those from healthy adult, the fitted 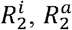, and *α* from neonates were substantially reduced but *ε*_0_ increased, consistent with incomplete myelination. Significant positive 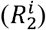 and negative (*α* and 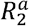) correlations were found with aging (demyelination) in elderly.

**Conclusion:** The developed framework can better characterize orientation dependences from a wide range of proton transverse relaxation measurements in human brain WM, shedding new light on myelin microstructural alterations at the molecular level.

## 1 INTRODUCTION

The intrinsic contrast of MR imaging of biological tissues comes mostly from spatial variations in proton relaxation rates,^1^ which are reflective of local environments and governed by varying molecular motions on different timescales.^2, 3^ In principle, proton relaxation measurements can provide basic microstructural information particularly for tissues with highly organized microarchitectures. For instance, myelinated axons in the human brain white matter (WM) are anisotropic and inhomogeneous in nature and water proton longitudinal (i.e., *R*_1_=1*/T*_1_) and transverse (i.e., *R*_2_=1*/T*_2_) relaxation rates have been revealed depending on orientations of axon fibers although the reported *R*_1_ relaxation anisotropy at 3T is much smaller than that of *R*_2_.^4-7^

Based on water proton transverse magnetization dephasing induced by applied directional diffusion gradients, diffusion tensor imaging (DTI) can provide axon orientation information in WM at an image voxel size level.^8, 9^ Generally, three orthogonal translational diffusivities (i.e., eigenvalues) and the corresponding directions (i.e., eigenvectors) relative to an external static magnetic field *B*_0_ can be determined based on the standard DTI model. While the magnitude of anisotropic diffusion can be well defined from the diffusion tensor eigenvalues, it remains unclear whether the direction of anisotropic diffusion can accurately represent an axon fiber direction, which is usually assumed as the direction of principal diffusivity.^9, 10^ When the direction of principal diffusivity was used as an internal orientation gold standard, some discrepancies appeared when compared with an axon orientation derived from either susceptibility tensor imaging^11-14^ or nanostructure-specific X-ray tomography.^15^ Similarly, when this internal reference was used to guide the orientation dependence of proton transverse (*R*_2_ and 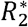) relaxation in vivo, the observed orientation dependence profiles manifested an angle offset *ε*_0_ that has not yet been accounted for.^4, 16-18^ Herein, *R* and 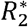 were treated equally regarding the axon fiber orientation dependences on *B*_0_.

In the past, susceptibility-based relaxation models have been proposed for characterizing anisotropic *R*_2_ and 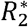 in WM,^14, 19-21^ expressed either by *A*_1_ + *A*_2_ cos 2*θ* + *A*_3_ cos 4*θ* or by *B*_1_ *B*_2_ sin^2^*θ* + *B*_3_ sin^4^*θ*. Here, *θ* is the angle between an axon fiber and *B*_0_, *A*_i_ and *B*_i_ (*i = 1,2,3*) the model parameters (i.e., trigonometric function coefficients). As revealed recently,^7, 18^ both functions are mathematically equivalent albeit with different coefficients. More importantly, these coefficients are not mutually independent as cos4*θ* (or sin^4^*θ*) can be expressed by cos2*θ* (or *sin*^2^*θ*) and vice versa; thus, any proposed biophysical interpretations of fitted model parameters will become ambiguous.

When considered as the potential origin of anisotropic *R*_2_ and 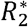 in WM, the magic angle effect (MAE) had not been appropriately evaluated in the literature.^20, 22^ Typically, the standard MAE function (i.e., orientation dependence) is written as *3cos*^2^*θ − 1*) ^2^, indicating that MAE will disappear with *θ*=54.7° (i.e., “magic angle”) and will be four times less with *θ* =90° than that with *θ* =0°. However, the observed *R*_2_ and 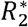 in WM contradicted this theoretical predication; in other words, the reported *R*_2_ and 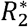 became larger when *θ* changed from 0° to 90°. As a result, MAE had been ruled out as a potential relaxation mechanism. It should be emphasized that this standard MAE function implicitly assumes that all restricted water molecules are uniformly orientated along the same direction, which might not be the case in brain WM.^23^ A general form of MAE function has long been available^24^ and lately reformatted to gain further insight into proton transverse relaxation orientation dependences.^25^

As both stemmed from thermally driven Brownian motions, molecular translational diffusion and rotational diffusion (or reorientation) should be intimately linked,^7, 26, 27^ with the former probed by DTI and the latter by MR relaxation measurements. To our knowledge, no direct connection exists between the two different measurements for studying the same anisotropic microstructures in WM.^28^ This disconnection might impede better characterization of rotationally restricted molecules both on the surface (water, H2O) and in the interior (lipid methylene, CH2)^29, 30^ of phospholipid bilayers in WM. In the past, an ultrashort transverse relaxation time 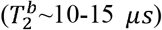 of semisolid lipid CH2 protons was observed by quantitative magnetization transfer (qMT) imaging,^31^ revealing the comparable orientation dependence with respect to that of surface water based on multiple orientation-dependent transverse relaxation studies in literature.^4, 7, 17, 18, 32^

The aim of this work was thus to introduce an angle offset *ε*_0_, determined by DTI diffusivities, into a cylindrical helix model based on the generalized MAE function for characterizing anisotropic transverse relaxation of ordered water and semisolid CH2 protons in WM. The proposed theoretical framework was validated by a high-resolution Connectome DTI dataset and then applied to previously published anisotropic *R*_2_ and 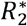 profiles at 3T in vivo from the human brain WM of neonates, healthy and diseased adults. The results demonstrate that the proposed model can better characterize the documented orientation-dependent proton transverse relaxation profiles.

## 2. THEORY

Water molecular reorientation in biological tissues, unlike in solution, becomes restricted to some extent,^33^ giving rise to a non-averaged dipolar interaction between two intramolecular proton nuclei, referred to herein as residual dipolar coupling (RDC) and signified by <H-H>. If the water rapidly rotates preferentially around a cylindrically symmetric axis that makes an angle *β* with <H-H> and an angle *θ* with *B*_0_, this RDC can be expressed by *DD * P*_2_ (*cos β*) *P*_2_ (*cos θ*), ^34, 35^ with the constant *DD* denoting the theoretical dipolar interaction strength for a rigidly fixed water and *P*_2_ (*x*) *=* (*3x*^2^ *−* 1)/2 the second order Legendre polynomial. P_2_ (cos *θ*) indicates the orientation dependence of RDC and ⟨ *P*_2_ (*cos β*) ⟩ is called “order parameter” in literature.^35, 36^ It has long been known^24^ that water proton signal splitting in partially hydrated tissues is proportional to the mean of RDC, i.e., ⟨(*P*_2_ (*cos θ*)⟩, and the corresponding linewidth or *R*_2_ relaxation rate in fully hydrated tissues depends on the variance of RDC, i.e., ⟨ *P*_2_ (*cos θ*))^2^⟩. Note, the angle brackets stand for an ensemble or time average.

In the human brain WM, water molecules close to the hydrophilic surface of phospholipid bilayers, as schematically shown in Figure 1A, are highly organized as revealed by an advanced Raman scattering microscopic imaging study.^23^ Specifically, the direction of water RDC is parallel to the bilayer surface (Fig. 1B) or perpendicular to the surface normal (green dashed line). As the lipid’s long hydrocarbon chains are predominantly aligned with and rapidly rotate around the surface normal (Fig. 1C),^29, 37^ the direction of average RDC from lipid’s methylene (CH2) groups becomes orthogonal to that from water in WM (Fig. 1B). It is worth mentioning that these *static* RDC configurations are only valid on the timescale of 10^−14^ s, i.e., vibrational frequencies from the pertinent chemical bond stretching modes^23^ that are unrelated to proton MR relaxation.^2, 3^

**FIGURE 1.**
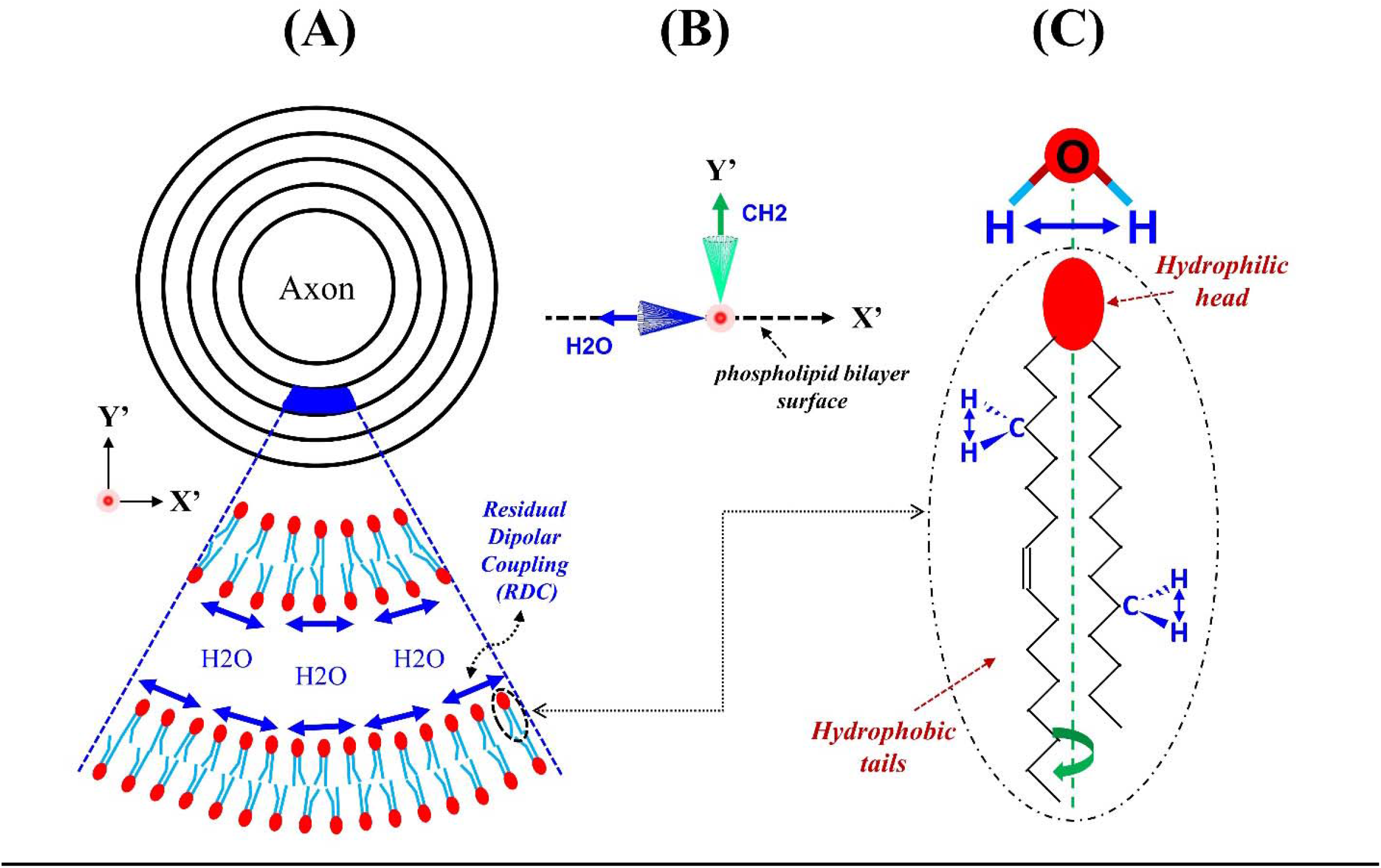
Schematics of residual dipolar coupling (RDC, blue double-headed arrows) organized around an axon fiber (A). A lipid molecule is highlighted (C) with RDCs (B) of ordered water and methylene (CH2) protons respectively perpendicular to and parallel to bilayer surface normal (green dashed line). Figures 1A and 1C were adapted from Figure 2 in the reference.^23^

As demonstrated in Figure 2A, a concentric distribution of RDCs around an axon can be rearranged into a specific cylindrically symmetric system with an open angle *α =* 90*°*. In an imaging voxel from biological tissues, another extreme case with *α* ≈ *0°* (Figure 2B) could coexist as well in which all RDCs are uniformly orientated along one direction.^38, 39^ A general function of orientation-dependent *R*_2_ has been obtained by averaging (*3 cos*^2^*θ* − 1)^2^ in an axially symmetric system.^24, 25^ Equations 1 and 2 provide respectively the reformatted^25^ and the original^24^ functions with pertinent angles defined as before and depicted in Figure 2C. The angle *θ*, formed between <H-H> and *B*_0_, can be expressed by three different angles *α, ε* and *φ* denoting respectively between <H-H> and the symmetry axis 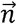, between 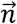 and *B*_0_, and the azimuthal angle of <H-H>. An ensemble or time average must be taken over all *φ*. A few representative orientation (*ε*) dependences of *R*_2_ with different *α* are exemplified in Figure 2D.

**FIGURE 2.**
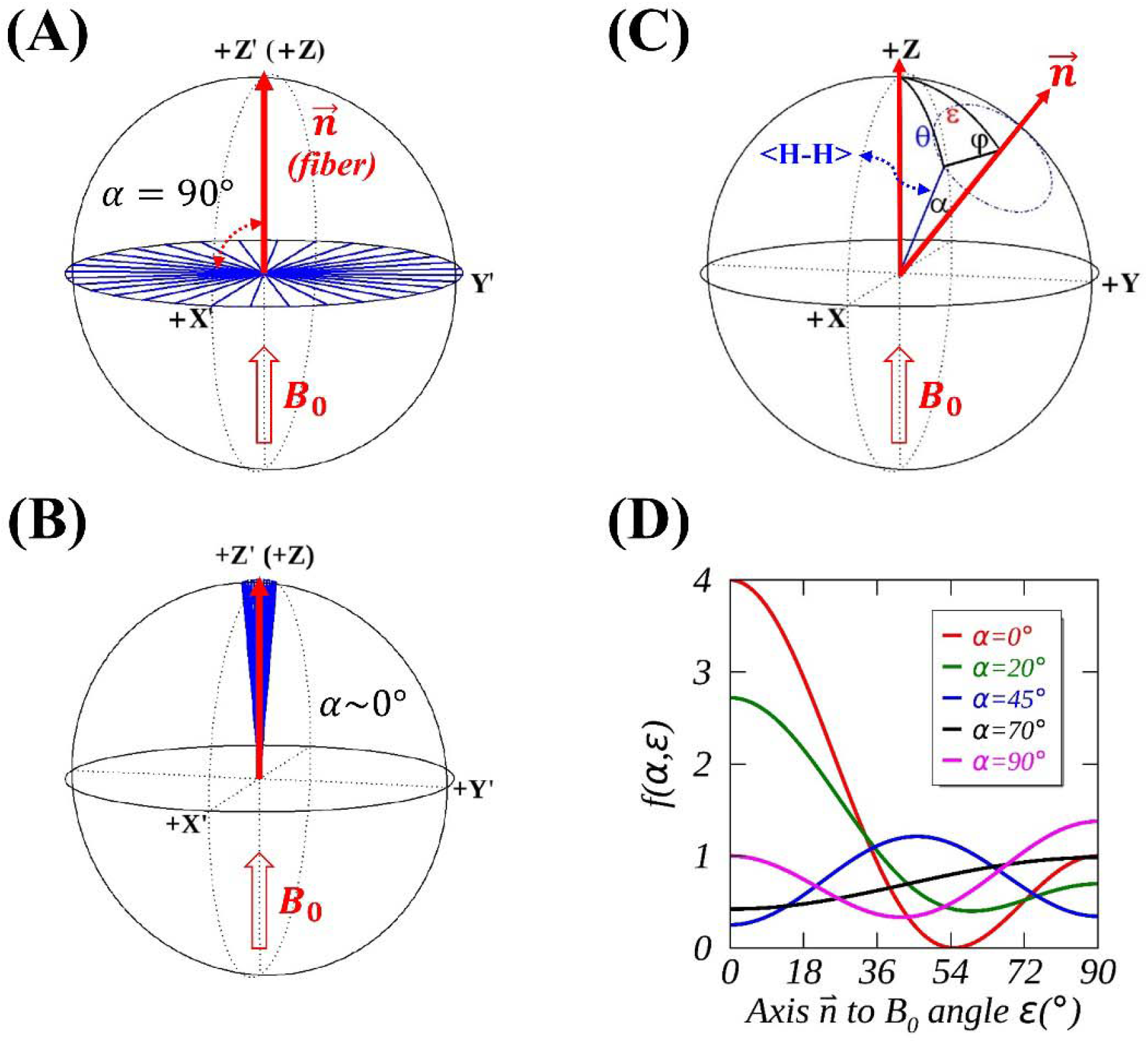
An axially symmetric model with an open angle *α*=90° (A) or *α∼*0*°* (B) when *ε* =0°, i.e., the primary axis 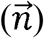 of an axon fiber aligned with *B*_0_. A general case with *α* ≠ 0*°* and *ε* ≠0*°* is also presented (C) and five anisotropic *R*_2_ orientation dependence *f* (*α, ε)* profiles are plotted (D) with *α* ranging from 0° (red) to 90° (magenta).

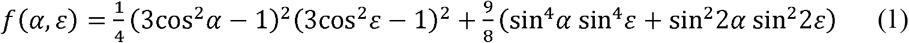

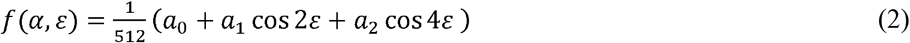

Here, the coefficients *a*_0_, *a*_1_ and *a*_2_ in Eq. 2 are respectively defined by 81cos4*α*+156cos2*α*+467, 180cos4*α*+432cos2*α*+156, and 315cos4*α*+180cos2*α*+81 as originally presented by Berendsen.^24^ It should be mentioned that Eq. 2 was previously employed for characterizing both isotropic and anisotropic transverse relaxation contributions. In this work, however, *f* (*α, ε*) was used only for modeling anisotropic transverse relaxation.

The orientation information can be obtained by physically rotating WM specimens relative to *B*_0_ for studies ex vivo.^19, 20, 40^ Because the first measurement may not start at *ε* =0°, an angle offset *ε* _0_ has to be introduced into Eq. 1 as recently demonstrated.^41^ For studies in vivo, on the other hand, the orientation information must be inferred from DTI.^16, 42^ Specifically, relative to *B*_0_, the primary eigenvalue direction of diffusion tensor is *assumed* the predominant orientation of axon bundles within an image voxel.^8, 9^ If this assumption is valid, the theoretical *R*_2_ or 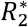 orientation dependence profiles will not be offset whereas the prior measurements demonstrated otherwise.^17, 18, 31, 43^ Therefore, *ε* _0_ has been incorporated in Eq. 1 for studies in vivo in the current work.

In addition, *ε*_0_ was hypothesized linking directly to the direction of a complete anisotropic translational diffusion. For a typical “zeppelin” diffusion tensor, neither axial (*D*_‖_) nor radial (*D*_⊥_) diffusivity characterizes such an anisotropic diffusion. For convenience, *D*_. ⊥_ was assumed arising solely from an isotropic diffusion, and the direction of the vector difference between *D*_‖_ and *D*_⊥_ could be considered along 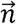 as depicted in Figure 3B, thus resulting in an angle offset *ε* _0_ when *D*_‖_ was considered as a reference. If average *D*_‖_ and *D*_⊥_ are available from whole WM, an average *ε*_0_ can be calculated by tan^-1^(*D*_⊥_ */D*_‖_) and compared with the fitted *ε*_0_ to validate the hypothesis. Even better when voxel-based *ε*_0_ is available and has been accounted for, the measured transverse relaxation orientation dependence should be consistent with the theory, without an angle offset *ε*_0_, if the hypothesis is valid.

**FIGURE 3.**
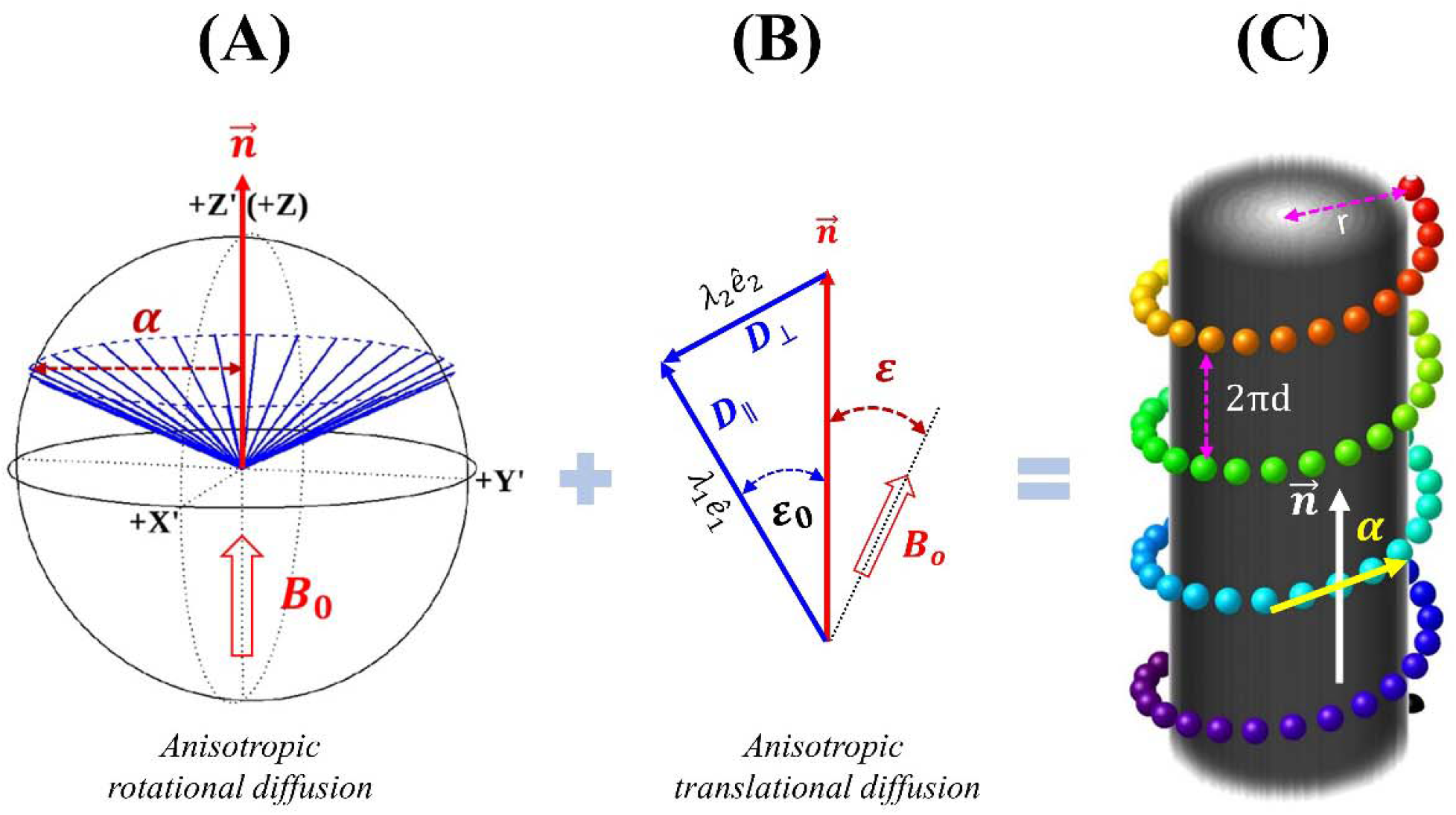
A combined anisotropic rotational diffusion (A) and anisotropic translational diffusion (B) into a cylindrical helix model (C), with an open angle *α* and circular helix parameters (i.e., *r*, radius and *2πd*, pitch) linked by tan *α = r/d*. An angle offset (*ε*_0_) is determined by directional diffusivities (*D*_‖_ and *D*_⊥_).

To conceptually connect an anisotropic rotational motion with an anisotropic translational diffusion, a standard axially symmetric system (Figure 3A) could be transformed into a cylindrical helix model (Figure 3C). Although sharing the same geometric microstructure, the proposed model is radically different from the previously developed hollow cylinder fiber model (HCFM)^44-46^, with the former derived from intramolecular residual dipolar interactions but the latter based on the “susceptibility effect”. Further, the proposed model deals exclusively with anisotropic *R*_2_ originating from both within (Fig. 2A) and outside (Fig. 2B) myelin sheaths. When the complete anisotropic translational diffusion is incorporated, the proposed cylindrical helix model has captured the most relevant degrees of molecular motions responsible for the observed shifted orientation-dependent *R*_2_ profiles in WM.

Mathematically, an open angle *α* and circular helix parameters, i.e., radius (*r*) and pitch (2*πd*), become connected via the relationship of tan *α = r/d*. Herein, *r* could be considered as a sum of an average thickness of myelin sheaths and an average nonmyelinated axon radius. Note, the directions of both translational and rotational molecular motions are considered collinear, pointing to an axon primary axis. Given a fixed length of helix pitch, the helix radius will asymptotically grows as *α* becomes widened. On the other hand, if a water molecule had been spiraling upward around a typical axon, with translational diffusion coefficient *D*_*a*_ and rotational correlation time *τ*_*b*_, the helix pitch would have been equal to 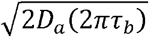.

## 3 METHODS

Unless otherwise specified, all orientation-dependent transverse relaxation rate profiles, identified by letters in an alphabetical order in Table 1, were extracted from image-based graphs in previous publications using a free online tool (www.graphreader.com), and replotted in the figures (black triangles) herein. The orientations (Φ) associated with the measured *R*_2_ and 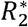 were originally determined by the primary diffusivity direction 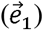 from DTI relative to the main magnetic field 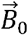, i.e., 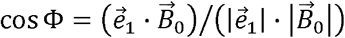, with Φ *=ε* + *ε* _0_ as depicted in Figure 3B. These estimated Φ values were sorted by increasing angles from 0*°* to 90°, and further averaged within a predefined interval or angular resolution (AR). The highest *b* values used in DTI were also provided in Table 1. *R*_2_ and 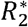 mappings were acquired using conventional Carr-Purcell-Meiboom-Gill (CPMG) spin-echo and gradient-echo sequences, respectively. The original relaxation data were acquired from whole brain WM in vivo on 3T MR systems following the ethical guidelines as stated in the original publications; but the use of these extracted data was not reconsented. If not stated otherwise, all concerned relaxation metrics were related to restricted water molecules in WM.

**TABLE 1.**
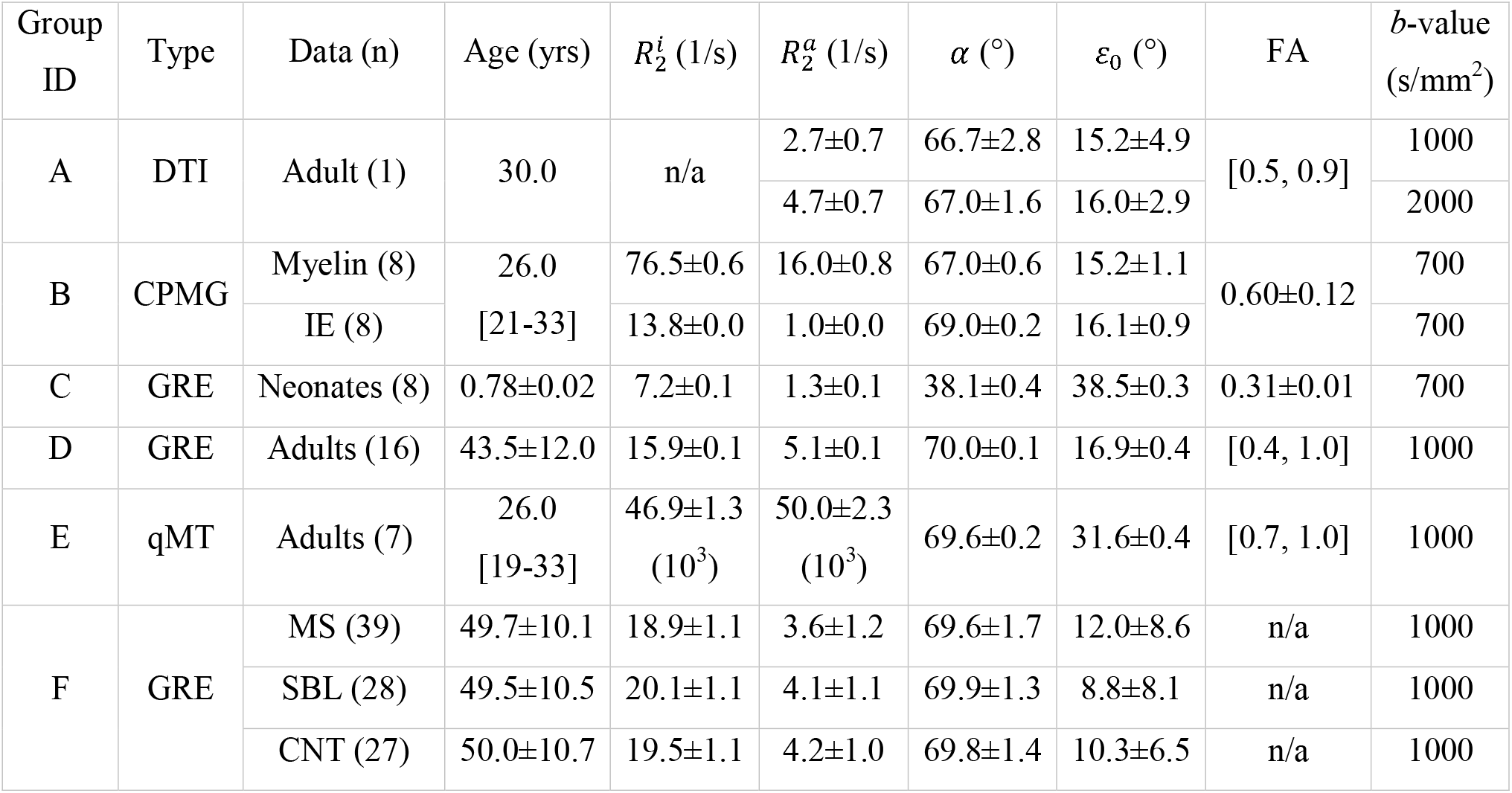
The fits from Fit A (based on Eq. 4) for previously published anisotropic *R*_2_ and 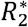 and related DTI metrics in WM in vivo at 3T. Data are denoted by means ± standard deviations or range (square brackets) if applicable.

### 3.1 Anisotropic *R*_2._ derived from Connectome DTI (Group A)

A public domain high-resolution (i.e., an isotropic voxel size of 760 *μm*^3^) Connectome DTI dataset of one healthy human brain (age=30 years)^47^ was utilized for validating the proposed theoretical framework regarding *ε*_0_. More specifically, paired with the associated *b*=0 (s/mm^2^) images, 6 preprocessed data subsets with *b*=1000 (s/mm^2^) and 12 with *b*=2000 (s/mm^2^) were individually analyzed using FSL DTIFIT^48^ to generate the following fit parameters: eigenvalues (*λ*_*i*_) and eigenvectors 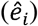, with *i* = 1, 2, 3; fractional anisotropy (FA); mode of anisotropy (MO);^49^ and T2W signals (SO) with *b*=0 (s/mm^2^). Supplementary Figure S1 provides exemplary fits for a rectangular ROI (in black) from corpus callosum as pointed by a red arrow in Figure 4D. The mean of the fits from data subgroup with *b*=1000 (s/mm^2^) or *b*=2000 (s/mm^2^) were used independently for further analysis.

**FIGURE 4.**
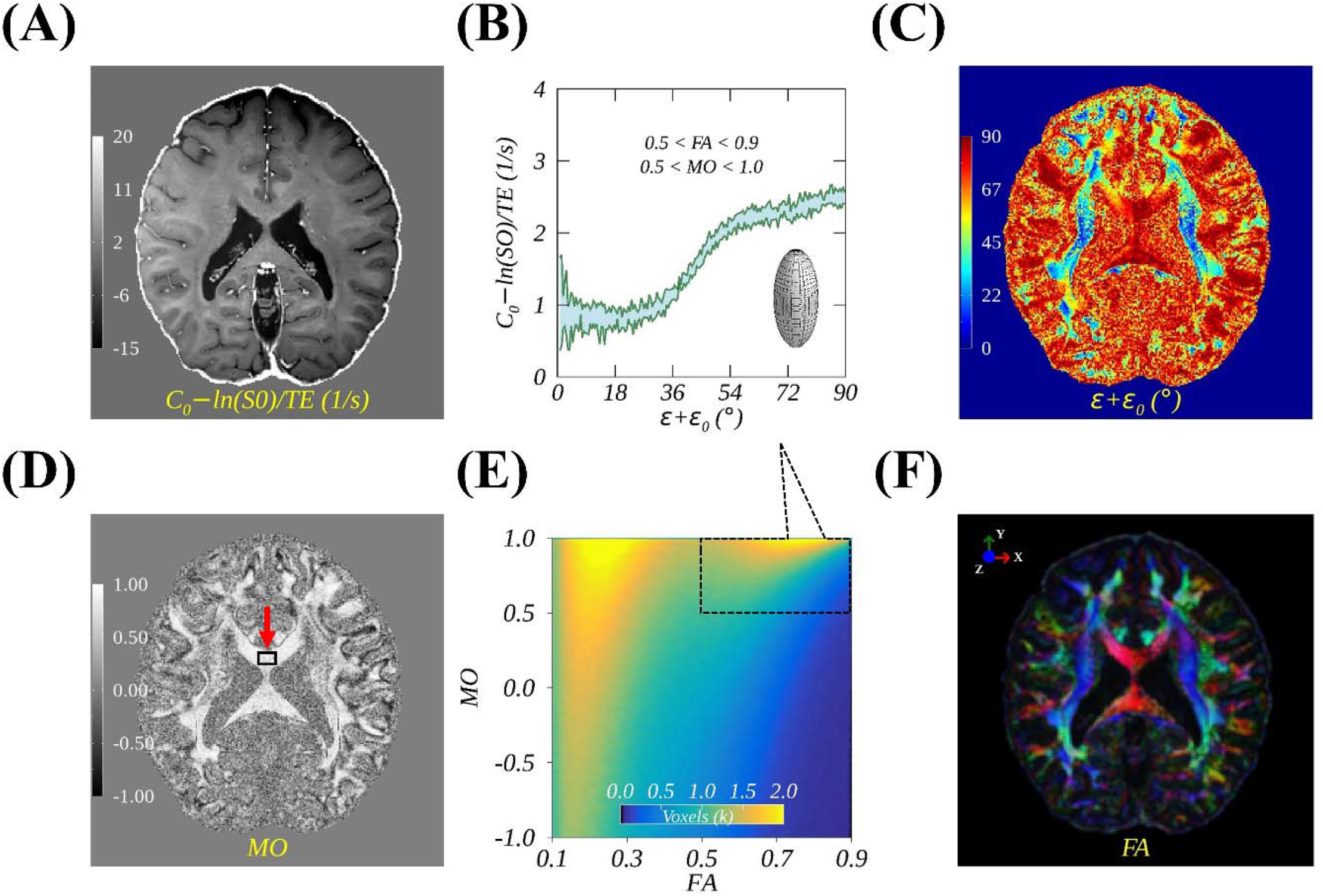
Average DTI parametric maps of anisotropic *R*_2_ = *C*_0_ − ln (*SO*) /*TE* (A), directions of principal diffusivities (C), modes of anisotropies (D), and colored fractional anisotropies (F). An anisotropic *R*_2_ orientation dependence profile (mean ± SD) (B) is shown for WM voxels defined (rectangular box) by limited FA and MO ranges (E) in whole brain. These DTI fits were derived from diffusion weighting data subsets (n=6) with *b*=0, 1000 s/mm^2^, which are publicly available.^47^

Given a known TE=75ms used in DTI,^47^ anisotropic *R*_2_ (denoted by 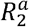) could be readily assessed from T2W signal based on Eq. 3 as demonstrated previously.^50^ A similar idea has been exploited lately for separating an isotropic 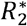 component from a single GRE measurement.^51^

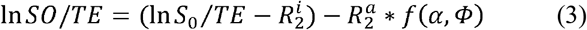

Note, SO became *S*_0_ (i.e., apparent proton density) when TE=0 and the term (ln 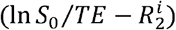) was treated as a constant *C*_0_ to be determined by fitting. Both *S*_0_ and 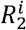 (i.e., isotropic *R*_2_ component) were reportedly least varied in WM.^4^ Within the whole brain, the measured SO values (in a logarithmic scale) from specific voxels possessing a linear (i.e., 0.5<FA<0.9 and 0.5<MO<1.0) diffusion anisotropy were sorted based on the calculated fiber orientations (Φ) and then averaged into 180 different bins ranging from 0° to 90°, i.e., AR=0.5°.

### 3.2 Compartmental anisotropic *R*_2._ from myelin water imaging (Group B)

Anisotropic *R*_2_ orientation dependence profiles of myelin water and intra- and extra-cellular (IE) water were retrieved respectively from Figures 7A and 7B in a recent study of orientation-dependent myelin water imaging.^18^ In this work, Figures 6A and 6B replicate the orientation-resolved (AR=5°) compartmental *R*_2_ values, measured from eight healthy volunteers with a mean age of 26 years (range=21-33 years). An interval between two adjacent refocusing pulses in CPMG pulse sequence was 8 ms. The highest *b*-value used in DTI was 700 s/mm^2^.

**FIGURE 5.**
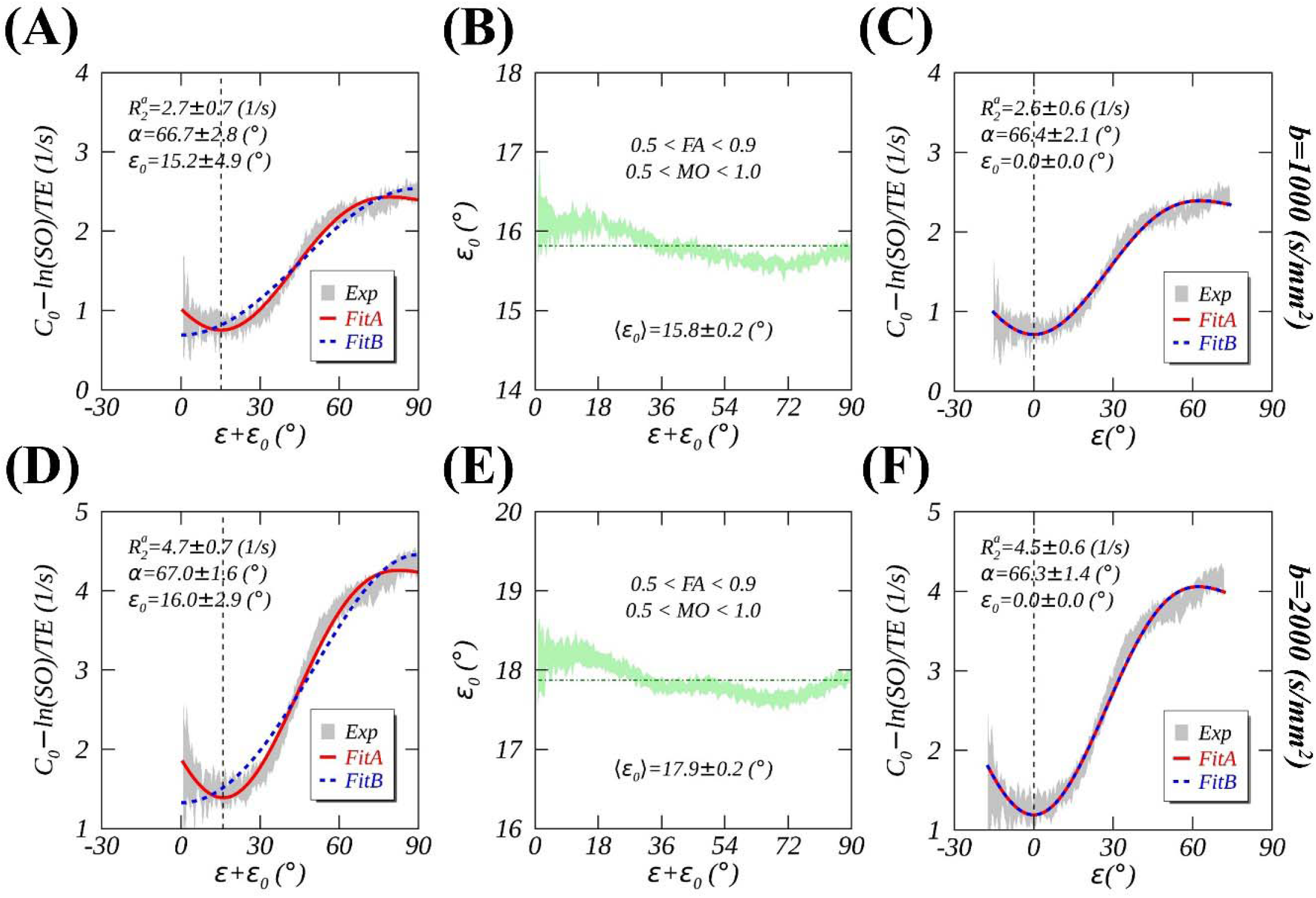
Demonstrations of angle offsets *ε*_0_ depending on directional diffusivities. Anisotropic *R*_2_ orientation dependences characterized by Fit A (red solid lines) and Fit B (blue dashed lines) before (A and D) and after (C and F) correcting angle offsets *ε*_0_ for data with *b*=1000 s/mm^2^ (A-C) and *b*=2000 s/mm^2^ (D-F). Orientation dependences of *ε*_0_ are also presented (B and E). These anisotropic *R*_2_ profiles were generated from publicly available Connectome DTI datasets.^47^

**FIGURE 6.**
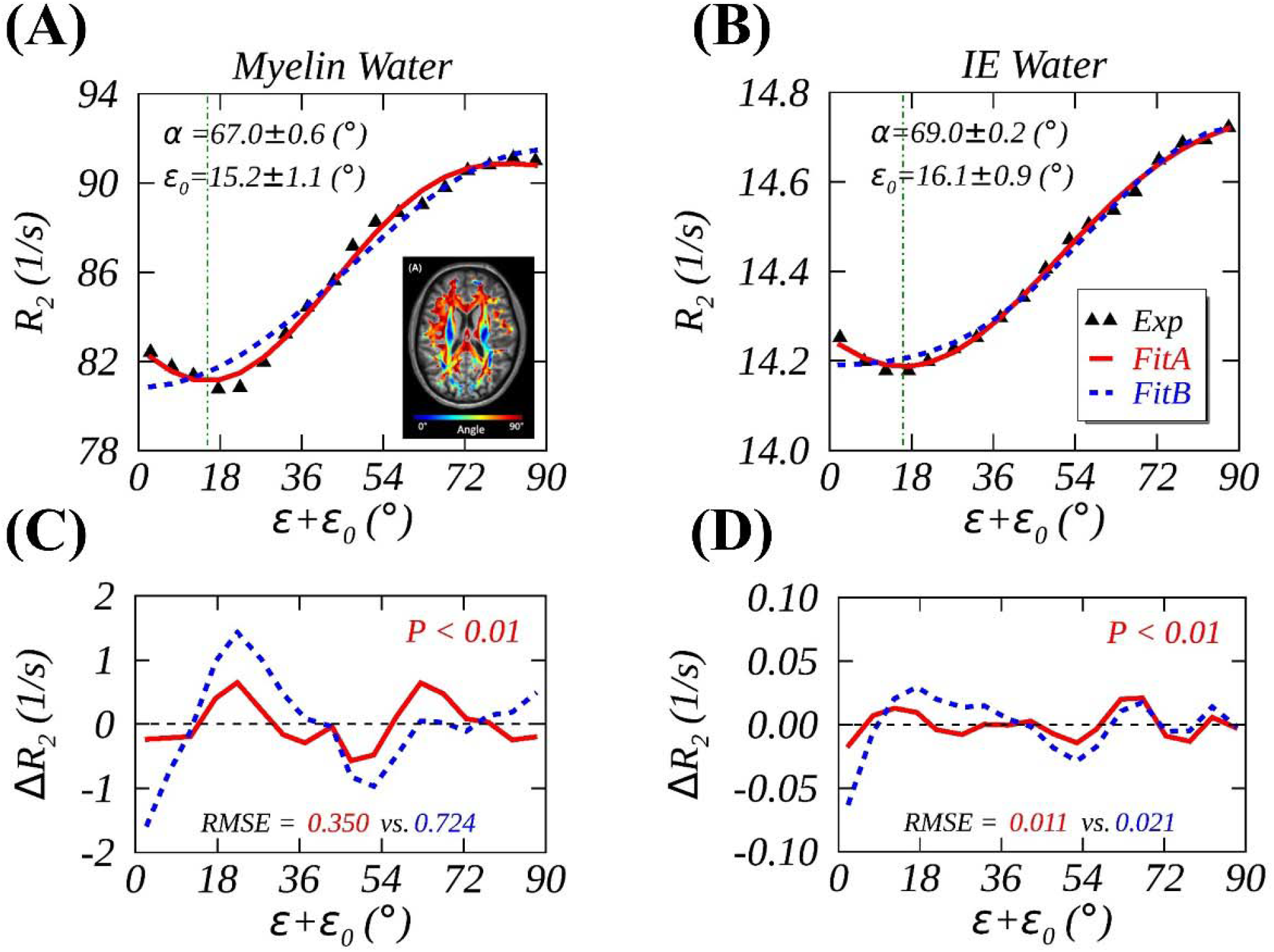
Measured (black triangles) and fitted (Fit A, red solid lines; Fit B, blue dashed lines) anisotropic *R*_2_ of myelin water (A) and intra- and extracellular (IE) water (B) in human brain WM at 3T in vivo, with fitting residues Δ*R*_2_=Fitted-Measured (C and D). The measured data and an inset image (A) were retrieved and adapted from the reference.^18^

**FIGURE 7.**
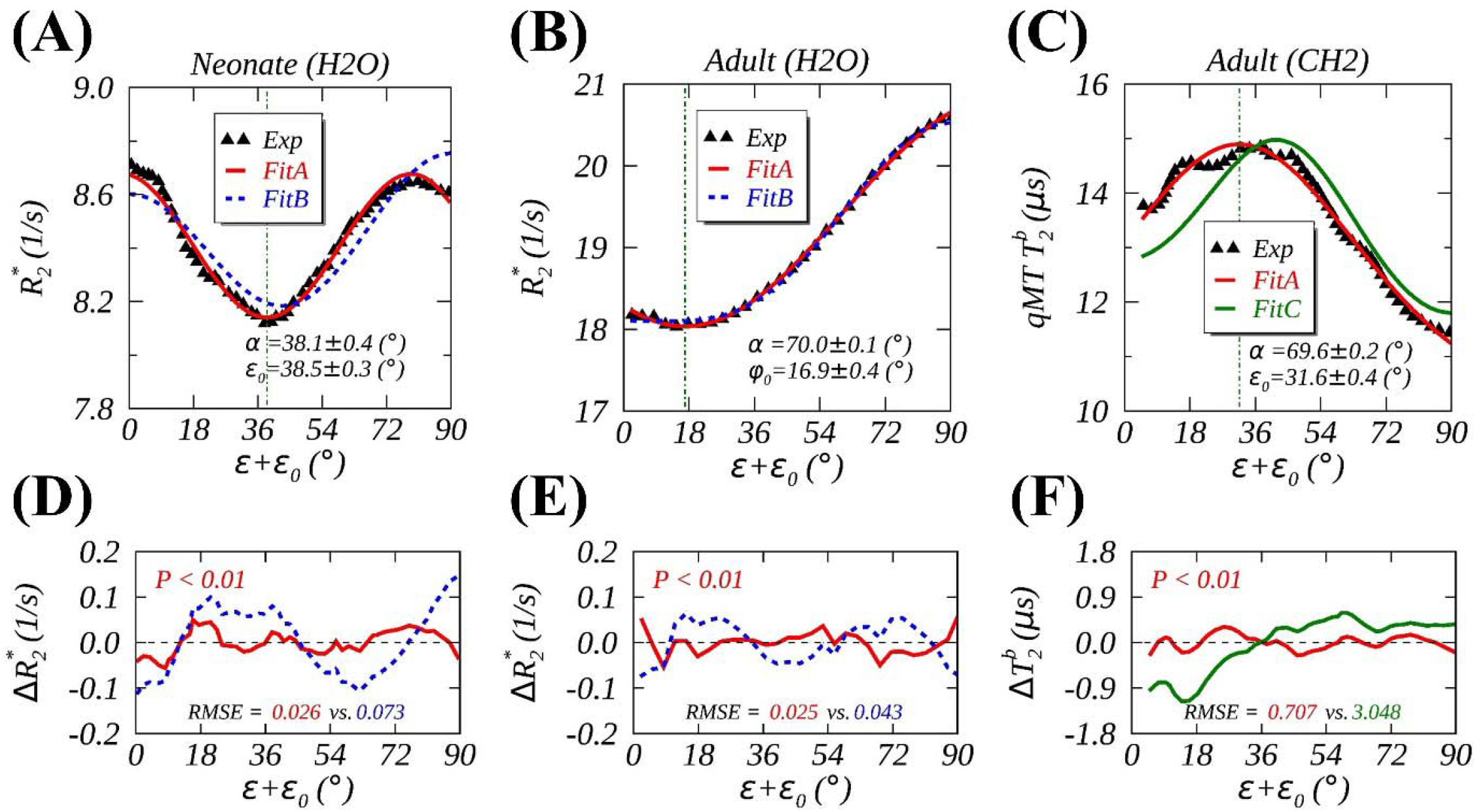
Measured (black triangles) and fitted (Fit A, red solid lines; Fit B, blue dashed lines, Fit C, green solid lines) anisotropic 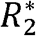 from ordered water in WM of neonates (A), healthy adults (B), and 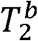 of semisolid macromolecules in WM of healthy adults (C) at 3T in vivo, with fitting residues Δ*R*_2_=Fitted-Measured (D-F). The measured data were respectively retrieved from three references.^4, 17, 31^

In another prior publication,^52^ average axial (*D*_‖_ =1.44±0.24 μm^2^/ms) and radial (*D*_⊥_ =0.47±0.11 μm /ms) diffusivities in WM were reported in Table 1 for an age-matched group at 3T. As defined in Figure 3B, an average *ε*_0_ and its uncertainty Δ*ε* _0_ were respectively calculated by tan^-1^(1/*ρ*) and Δ*ρ* /(1+*ρ*^2^), with *ρ* =*D*_‖_/*D*_⊥_ and Δ*ρ* derived from Δ*D*_‖_ and Δ*D*_⊥_ following basic error propagation rules.^53^ Note, *ρ* was listed as 3.33±1.29 (μm^2^/ms) in the original paper and used in the calculation in this work. Fractional anisotropy (FA) was also computed^8^ and tabulated in Table 1 herein.

### 3.3 Anisotropic 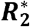 from neonates and adults (Groups C and D)

Orientation-dependent 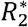 profile (AR=0.33°) in WM from 8 term neonates (mean age=40.4±1.1 weeks) was retrieved from Figure 3C in a previous study (Group C)^17^ and replotted in Figure 7A herein. The highest *b*-value used in DTI was 700 s/mm^2^. The directional diffusivities were also reported in the original paper, i.e., *D*_‖_ =1.69±0.03 (μm^2^/ms) and *D*_⊥_ =1.06±0.04 (μm^2^/ms). In a different study on adult WM,^4^ orientation-dependent 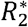 measurements (AR=3°) were performed on a group of 11 healthy female and 5 healthy male subjects with an average age of 43.5 ± 12 years (range: 26–67 years, median: 43 years). An average anisotropic 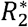 profile (age ≥ 43 years) was retrieved from Figure 2g in the original publication^4^ and reproduced in Figure 7B in this work. The highest *b*-value used in DTI was 1000 s/mm^2^.

### 3.4 Anisotropic 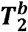 deduced from semisolid macromolecules in WM (Group E)

qMT imaging was performed on 7 healthy subjects (mean age=26 years, range=19–33 years).^31^ The system characteristic MT parameters were determined based on a two-compartment model comprising a liquid pool of free water and a semisolid pool of macromolecules. The derived average transverse relaxation time 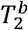 orientation dependence profiles of semisolid macromolecules in WM were retrieved from Figure 7a in the original paper and replicated in Figure 7C in this work. The highest *b*-value used in DTI was 1000 s/mm^2^. The reported proton 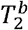 was largely associated with semisolid methylene (CH2) groups on lipid hydrocarbon chains in the interior of lipid bilayers in WM.^31^

### 3.5 Anisotropic 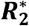 from people with MS and two controls (Group F)

A clinical study of anisotropic 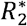 orientation dependences in WM was conducted among people with multiple sclerosis (MS, n=39; mean age=49.7±10.1 years), their age-matched asymptomatic siblings (SBL, n=28) and unrelated healthy controls (CNT, n=27).^32^ Anisotropic 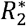 profiles (AR=5°) of each subject from three groups were retrieved from Supporting Information in the original paper, and group-average 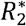 profiles were presented in Figure 8A herein. The highest *b*-value used in DTI was 1000 s/mm^2^.

**FIGURE 8.**
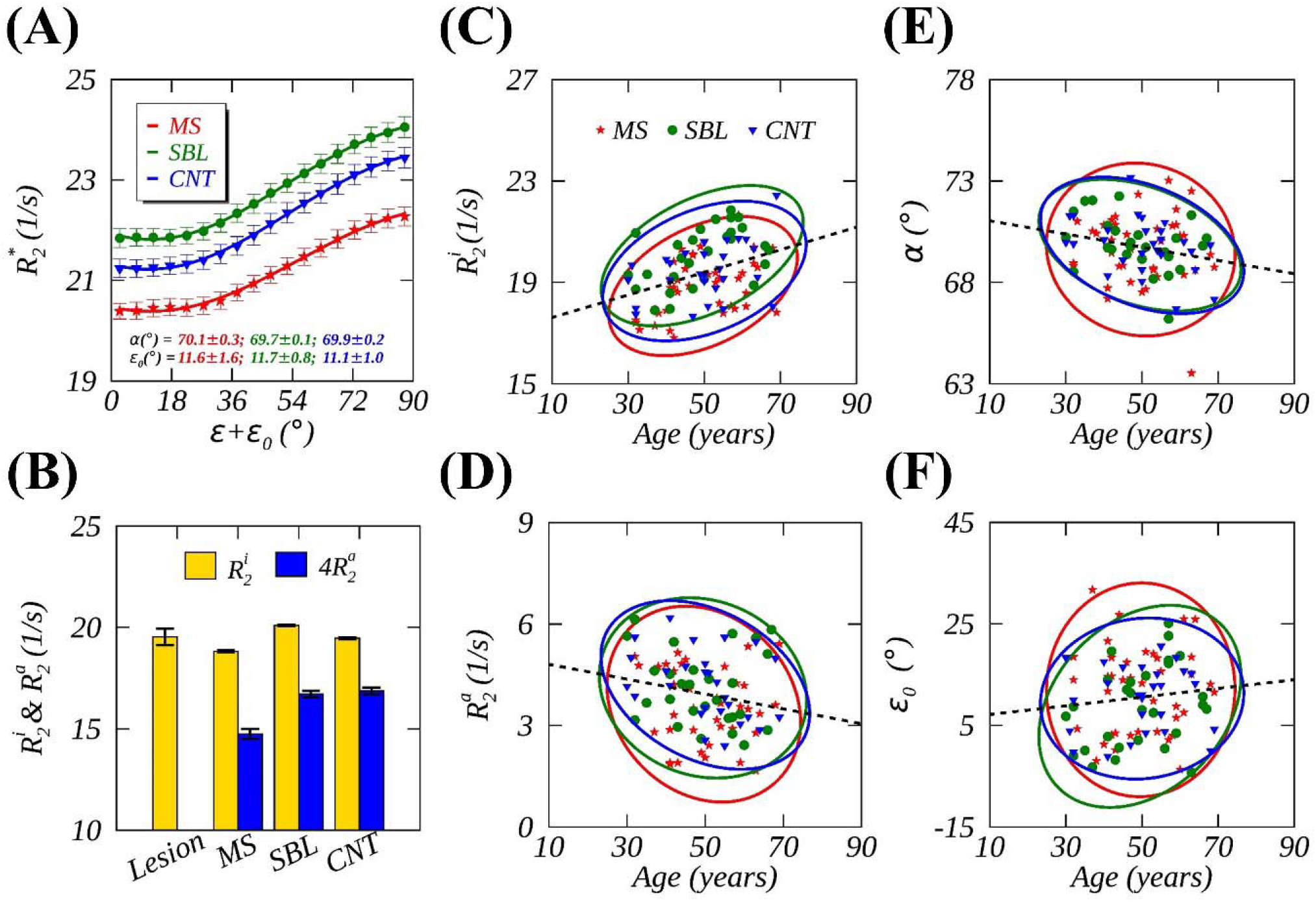
Measured (symbols) and fitted (Fit A, lines) anisotropic 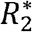 in WM (A) from people with multiple sclerosis (MS, red) and two controls (SBL, green; CNT, blue), with the fitted *α* and *ε*_0_ imprinted in colors, and with 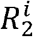 and 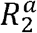 represented by color bars including those from lesions (B). Correlations between the fits from individual subjects and their ages are displayed for 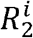 (C) 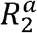 (D), *α* (E), and *ε*_0_ (F), overlaid with 95% confidence ellipses from individual subgroups and linear regression lines (black dashed lines) from combined subgroups. The measured 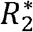 data were retrieved from Supporting Information in the reference.^32^

### 3.6 Nonlinear least-squares curve fittings

An optimization method of Levenberg-Marquardt nonlinear least-squares, implemented in an IDL script from the public domain (http://purl.com/net/mpfit),^54^ was used for modeling anisotropic *R*_2_ and 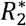 orientation dependence profiles, based on the proposed model (*R*_2,1_) in Eq. 4, herein referred to as “Fit A”.

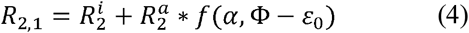

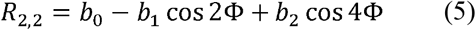

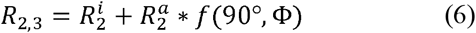

There were four model parameters and one independent variable (Φ) in Eq. 4, i.e., 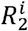 and 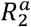 denoting orientation independent and dependent contributions, an open angle *α* for a particular RDC distribution, and an angle offset *ε*_0_. The same data from some Groups were also fitted using a previously developed model *R*_2,2_ (without *ε*_0_) as written in Eq. 5 (labeled as “Fit B”). It should be emphasized that when Fit B included *ε*_0_, it would become equivalent to Fit A albeit with different model parameters. With *α* fixed to 90° and *ε*_0_ set to 0°, *R*_2,1_ was transformed into *R*_2,3_ (Eq. 6), referred to as “Fit C”. For consistency, T2W signal fitting in Eq. 3 was categorized in the same way as in Eq. 4, i.e., referred to as either “Fit A” or “Fit B” with or without *ε*_0_ in *f* (*α*, Φ), respectively.

Initiating with five sets of different values within different constraints, the curve fittings were unweighted and constrained, ^55^ with the limited ranges for model parameters: *C*_0_ =[50, 200]; 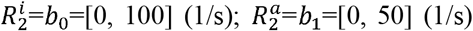; *α*=[0°, 90°]; *ε*_0_=[-45°, 45°]; and *b*_2_=[0, 10] (1/s). Note, 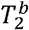 in units of *μs* from Group E was converted into 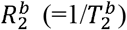 and then scaled down within the scale of fitting parameter constraints. Goodness of fit was characterized by root-mean-square error (RMSE), based on the measured (“Exp”) and fitted (“Fit”) transverse relaxation orientation dependence profiles (represented by *N* data points), i.e., 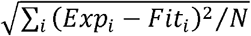, with *i* changing from 1 to *N*. The statistical significance was assessed by an *F*-test when comparing two fitting models. A *P*-value was derived from *F*-distributions with significance indicated by *P* ≤ 0.05.

All the reported fitting results (mean ± standard deviation) were obtained from Fit A unless specified otherwise. Pearson correlation coefficients (PCC) were listed in Table 2 between the fits from Group F and the age of each participant from individual or combined subgroups. The statistical significance of observed linear correlations was also set to *P* ≤ 0.05. All image and data analysis were performed using customized codes written in IDL 8.8 (Harris Geospatial Solutions, Broomfield, CO, USA).

**TABLE 2.** Pearson correlation coefficients (PCC) between the fits (Fit A based on Eq. 4) of anisotropic 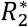^*^ in WM and the ages of people from individual or combined subgroups (Group F). MS, multiple sclerosis; SBL, age-matched asymptomatic siblings; CNT, age-matched unrelated healthy subjects.

| Fits | MS |  | SBL |  | CNT |  | ALL |  |
| --- | --- | --- | --- | --- | --- | --- | --- | --- |
|  | PCC | *P*-value | PCC | *P*-value | PCC | *P*-value | PCC | *P*-value |
| $R_2^i$ | 0.41 | 0.01 | 0.48 | 0.01 | 0.38 | 0.05 | 0.38 | <0.01 |
| $R_2^a$ | -0.19 | 0.25 | -0.13 | 0.51 | -0.34 | 0.08 | -0.20 | 0.05 |
| $\alpha$ | -0.06 | 0.72 | -0.37 | 0.05 | -0.38 | 0.05 | -0.23 | 0.03 |
| $\varepsilon_0$ | 0.01 | 0.96 | 0.30 | 0.12 | 0.07 | 0.75 | 0.11 | 0.28 |

## 4 RESULTS

### 4.1 Anisotropic *R*_2_. orientation dependence with an offset

Figure 4 presents four parametric maps derived from DTI (*b*=1000 s/mm^2^): (A) *C*_0_ − ln(*SO*)*/TE*, an equivalent of anisotropic *R*_2_, (C) orientations (i.e., Φ *= ε* + *ε*_0_) of principal diffusivities, (D) modes of anisotropies ranging from 1.0 to -0.1 denoting respectively an ideally linear (“stick”) and an ideally planar (“plate”) diffusion tensor, and (F) colored fractional anisotropies (i.e.,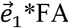). An anisotropic *R*_2_ orientation dependence profile was plotted (B) as a ribbon (width=mean ± SD, n=6). The image voxels from whole brain WM were selected according to limited FA and MO ranges as highlighted (black box) in 2D histogram (E). An ROI-based (pointed by a red arrow in Fig. 4D) average fits (mean ± SD) from all data subsets (n=18) for different parameters are plotted in Supplementary Figure S1 including three diffusivities (*λ*_*i*_, *i =* 1.2.3) and mean diffusivity (MD). Although derived from T2W images with a single TE, this characteristic profile was comparable to the previous in healthy adult human brains based on the standard *T*_2_ mapping with multiple varying TEs.

Figure 5 demonstrates the fitted anisotropic *R*_2_ profiles using Fit A (red solid lines) and Fit B (blue dashed lines) before (A and D) and after (C and F) correcting voxel-based angle offset *ε*_0_ derived from diffusivities with *b*=1000 s/mm^2^ (A-C) or *b*=2000 s/mm^2^ (D-F). Fit A significantly (*P*<0.01) outperformed Fit B without the voxel-based *ε*_0_ corrections; however, the differences between the two fits disappeared after correcting the voxel-based *ε*_0_. For data with *b*=1000 s/mm^2^, for instance, before and after the corrections. the calculated RMSEs between Fit A and Fit B were 0.081 vs. 0.137 and 0.081 vs. 0.081, respectively.

The voxel-based *ε*_0_ orientation dependences are profiled (mean ± SD) for DTI data with *b*=1000 s/mm^2^ (B) and *b*=2000 s/mm^2^ (E), showing that the fitted *ε*_0_ (vertical dashed lines, 15.2° ± 4.9°, A; 16.0° ± 2.9, D) and the calculated average ⟨*ε*_0_ ⟩ (horizontal dashed lines, 15.8° ± 0.2°, B; 17.9° ± 0.2°, E) become equivalent within the measurement errors. Although consistent with previous findings, the fitted 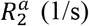 based on low *b*-value data (C) was substantially smaller (i.e., 2.6 ± 0.6 vs. 4.5 ± 0.6) than that with high *b*-value (F) on the same subject even after the *ε*_0_ corrections.

### 4.2 Anisotropic *R*_2_. from myelin water imaging

Figure 6 reveals that Fit A significantly (*P* < 0.01) outperformed Fit B for characterizing anisotropic *R*_2_ profiles (black triangles) of myelin water (A) and intra-and extra-cellular (IE) water (B) based on Group B data,^18^ indicated by reduced fitting residuals (i.e., Δ*R*_2_ =Fit-Exp, C and D). Quantitatively, the calculated RMSEs from Fit A, relative to those from Fit B, decreased almost half for myelin water (i.e., 0.350 vs. 0.724) and IE water (i.e., 0.011 vs. 0.021).

Between the two partitioned water pools, the fitted *α* (i.e., 67.0±0.6° vs. 69.0±0.2°) and *ε*_0_ (i.e., 15.2±1.1° vs. 16.1±0.9°) were comparable despite markedly different 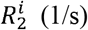(i.e., 76.3±0.6 vs. 13.8±0.0) and 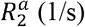 (i.e., 16.1±0.8 vs. 1.0±0.0). On the other hand, the calculated *ε*_0_ (i.e., 16.7±6.1°) based on average directional diffusivities in highly anisotropic (i.e., FA=0.60±0.12) WM was comparable with the fitted *ε*_0_ from anisotropic *R*_2_ profiles as expected.

### 4.3 Anisotropic 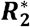 in WM from neonates and adults

Whereas both were better modeled using Fit A than using Fit B, the observed anisotropic 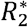 profile of neonates (Figure 7A) was substantially different from that of adults (Figure 7B). Specifically, the fits were smaller from neonates than from adults 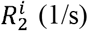, i.e., 7.2±0.1 vs. 15.9±0.1; 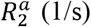, i.e., 1.3±0.1 vs. 5.1±0.1, and *α* (°), i.e., 38.1±0.4 vs. 70.0±0.1, but became larger for *ε*_0_ (°), i.e., 38.5±0.3 vs. 16.9±0.4 (vertical dashed green lines, A and B). The fitted *ε*_0_ (°) from neonates was slightly larger than the calculated counterpart (i.e., 38.5±0.3 vs. 32.1±1.1), probably due to an inaccurate replication of scattered 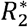 data from the original graph. Nonetheless, neonates possessed less anisotropic WM when compared with adults as indicated by the reported FAs (see Table 1). It is worth noting that a relatively reduced 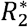 in neonatal WM stemmed largely from an anisotropic component and *R*_2_ was reportedly independent of *B*_0_ that is in good agreement with the proposed theoretical framework.^56^

### 4.4 Anisotropic 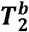 of semisolid CH2 in WM from adults

As demonstrated in Figures 7C and 7F, the profile of qMT-derived 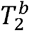 (i.e., 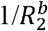) from image voxels with FA > 0.7 was significantly (*P* < 0.01) better modeled using Fit A than using Fit C. Despite originating from distinct proton groups and imaging methods, the fitted *α* (∼70°) appeared surprisingly close to that found in other adult groups as listed in Table 1. However, the fitted *ε* _0_ (°) was nearly twofold bigger than that from water proton 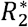 profile as shown in Figure 7B. Although consistent with the semisolid nature, the fitted 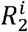 and 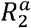 were unexpectedly higher (i.e., ∼50×10^3^ 1/s) when compared to the previously reported resonance linewidth of lipid chain protons from non-orientated bilayers, i.e., ∼3-6 kHz.^57^

### 4.5 Anisotropic 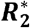 in WM with multiple sclerosis (MS)

Figure 8A shows average anisotropic 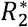 orientation dependence profiles for multiple sclerosis (MS, red), age-matched siblings (SBL, green) and unrelated healthy control (CNT, blue) groups. Although the fitted *α* and *ε*_0_ were comparable as shown by colored imprints (Figure 8A), the fitted 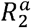 became the lowest for MS subgroup (Figure 8B) and so was 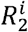 but to less degree. Figures 8C-F display scatterplots between individual ages and the corresponding fits of 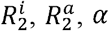, and *ε*_0_ from three subgroups, overlaid respectively with a 95% confidence ellipse.

Compared to those from two controls, 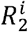 and 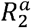 from MS subgroup were reduced but *ε*_0_ increased. Linear regression lines from combined subgroups were also included, highlighting significant (*P* ≤ 0.05) positive (PCC = 0.38 for 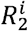) and negative (PCC = -0.20 and -0.23 for 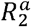 and *α*, respectively) correlations, as tabulated in Table 2, between aging (or demyelination) and the fits. The fitted *ε*_0_ more or less (PCC = 0.11, *P*=0.28) followed the similar trend of 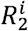. In general, an older people most likely possessed increased 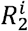 and *ε*_0_ but decreased 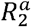 and *α*, in accordance with what had been known in neonates with developing WM (Figure 7A).

## 5 DISCUSSION

### 5.1 A cylindrical helix model for molecular rotation and translation

This work proposed a unique cylindrical helix model for characterizing anisotropic proton transverse relaxation orientation dependences in the human brain WM at 3T. This model shares some remarkable similarities with its predecessors.^14, 20, 21, 31^ First, the ultrastructure of myelin is described as multiple layers of lipid bilayers spiraling around an axon and further considered as the source of orientation-dependent transverse relaxation. Second, the most relevant direction is the orientation of an axon fiber, rather than individual directions of molecular constituents distributed around the axon. Third, other macromolecules such as membrane proteins in bilayers are ignored. Nevertheless, the proposed model possesses several unusual characteristics with respect to prior models.

First, it brings together anisotropic *R*_2_ and DTI diffusivities by an angle offset *ε*_0_. To our knowledge, this is the first time to combine the most relevant degrees of molecular motions for characterizing highly organized microstructures in WM.^26, 28^ Second, this proposed model is a generalized MAE model,^24, 25^ applied to not only ordered “liquid-like” water on the surface but also dynamic “solid-like” CH2 protons in the interior of bilayers. Further, it can be also used in peripheral nervous system, not to mention other tissues,^25^ where ordered water seemed uniformly (*α* ≈ 0*°*) aligned in elongated collagenous tissues as shown in Supplementary Figure S2.^58^ Third, this generalized model is built upon residual dipolar interactions,^24, 25^ fundamentally different from those based on tensorial magnetic susceptibility. Theoretically,^14^ the susceptibility-induced proton transverse relaxation rate is quadratically scaled with the strength of *B*_0_; as a result, this relaxation pathway will become more relevant only at a higher *B*_0_. As shown in Supplementary Figure S3, the susceptibility-induced contribution to the total relaxation rate was about 5% at 3T, based on *B*_0_ dependent *R*_1*ρ*_ dispersion data on rat brain.^59^ This theoretical prediction has been corroborated by the previous in vivo study showing that *R*_2_ from neonatal brain WM was principally independent of *B*_0_ ranging from 1.0T to 3.0T. These theoretical and experimental results have justified the relaxation mechanism underlying the proposed model, in excellent agreement with the previous finding in knee articular cartilage.^36, 60^

### 5.2 An angle offset *ε*_0_ depending on directional diffusivities

Initially, an angle offset _0_ was introduced as an ansatz for better modeling *R*_2_ orientation dependences in WM.^41^ Based on a high resolution Connectome DTI dataset,^47^ we provided strong evidence that *ε*_0_ is indeed modulated by DTI diffusivities although the detailed mathematical functions for related anisotropic molecular motions need to be constructed. As shown in Figure 5, the previously reported *R*_2_ would have relatively larger measurement uncertainties when close to “zero” degrees due to the low number of voxels possessing such orientations. To throw away these valuable yet imprecise data points would not be good practice as it could conceal the critical information underlying the actual biophysical mechanism.^43^

The fitted average ⟨ *ε*_0_ ⟩ and 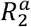 as shown in Figure 5 were larger when using a higher *b*-value in DTI. This result can be ascribed to decreased FA, stemmed mostly from reduced *D*_‖_ for a zeppelin tensor as demonstrated in Supplementary Figure S1. Therefore, ⟨ *ε*_0_ ⟩ became larger based on the relationship tan *ε*_0_=*D*_⊥_*/D*_‖_ (see Figure 3B). Because of the fixed voxel selection criterion (i.e., 0.5 < FA < 0.9) regardless of *b*-values, more image voxels with relatively higher FA were included in constructing *R*_2_ profiles with higher *b*-values, resulting in an enhanced 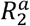 relative to that with lower *b*-values as demonstrated. These results suggest that some caution should be exercised when comparing previously reported *R*_2_ orientation dependence profiles. This important point has been clearly demonstrated in Figure 5 in the literature.^31^

Nonetheless, the fitted *ε*_0_ from previous transverse relaxation orientation dependences showed an increasing trend with aging in elderly (Figure 8F), consistent with an increasing *ε*_0_ from healthy adults to neonates (Figures 7A and 7B). These findings imply that *ε*_0_, or *D*_⊥_*/D*_‖_, appeared to be modulated by an extent of myelination in WM, in good agreement with the literature in which an increased myelination was linked to a decreased *D*_⊥_. ^61^ In an extreme scenario, *ε*_0_ can be zero if anisotropic *R*_2_ is induced exclusively by oriented water on the surface (i.e., *α*=90°) of lipid bilayers. With the absence of myelin in newborns,^62^ *ε*_0_ can also become zero in WM, corresponding to an ideal intra-axonal compartment with *D*_⊥_ normally set to zero.^7,27^

The fitted *ε* _0_ from anisotropic 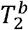 of semisolid CH2 protons (Figure 7C) deserves further explanation. The reported 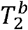 orientation dependence was guided by the direction inferred from the diffusion of water rather than lipid molecules. Moreover, lipid hydrocarbon chains may not be perfectly orientated along the bilayer normal as assumed (Figure 1B) particularly for more complex membranes that are rich in cholesterol (such as myelin).^63^ Hence, it is not surprising that the discrepancy appeared in the fitted *ε*_0_ as shown between Figures 7B and 7C.

### 5.3 An angle *α* for specific RDC distributions

A general form of MAE function *f* (*α, ε*) as written in Eq. 1 can be thought as a combination of two extreme cases, i.e., *f (*0*°,ε*) and *f* (90*°,ε*), with different weighting indicated by an open angle *α*. If observed anisotropic transverse relaxation in WM had resulted exclusively from ordered water within myelin multiple layers, the fitted *α* would have become 90° as shown in Figure 2A. The actual fitted *α* was, however, close to 70° for adult brain WM based on either ordered water or semisolid CH2 proton *R*_2_ relaxation measurements. This result implies that some uniformly aligned (i.e., *α*=0°) RDCs, at least from ordered water, had made considerably contributions. Previously published studies showed that these contributions increased as myelination decreased in WM of neonates^17^ until they completely dominated when myelin was absent in WM of newborns.^62^ This argument might help us to understand the reported increased *R*_2_ anisotropy in the very preterm infant brain when compared with that from the late preterm infant brain.^64^

For spins on lipid chains, *D*_*a*_ =4*10^−3^ (*μ m* ^2^/ms) and *τ*_*b*_ =100 (ms/rad) were reported^65^ and *d*=0.36*μm* could be determined based on the proposed model (Figure 3C) using the equation 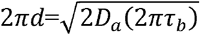. Further, an open angle of *α* could be determined to be approximately 70°, close to the findings from this work, if the radius of a representative axon is assumed 1.0 *μm* ^66^ To match the same *α* for restricted water, *D*_*a*_ =2.0 (*μm*^2^/ms)^27^ and *τ*_*b*_ =0.2 (ms/rad) ought to be expected. Whereas the value of *D*_*a*_ for restricted water seems not unreasonable in WM in vivo, the value of *τ*_*b*_ is uncertain.^46, 67^ More research is thus warranted.

### 5.4 Isotropic *R*_2_ relaxation contributions

According to Eq. 4, the observed proton MR transverse relaxation rates were divided into isotropic 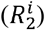 and anisotropic 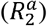 components, essentially assuming fast spin exchange between two environments.^68^ The former may arise from several relaxation pathways such as intrinsic dipolar interactions modulated by fast (e.g., pico- and nanoseconds) timescales of molecular motions,^2, 50^ chemical exchange and diffusion due to slow (e.g., micro- and milliseconds) molecular motions,^46^ and non-local magnetic susceptibility inhomogeneity 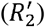 if gradient echo rather than spin echo signal was used.^20^ Contrast to the intrinsic dipolar interactions on fast timescales, the effect of slow molecular motions on 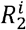 can be suppressed by *R*_1*ρ*_ dispersion.^46, 69^

The fitted 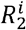 was reduced in developing (Figure 7A) and pathological (Figure 8C) WM compared to that in the healthy brain, consistent with fewer macromolecular labile protons. It was also shown^41^ that 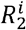 was considerably larger at ≥ 7T ex vivo when compared to those at 3T in vivo, possibly due to an increased *B*_0_ inhomogeneity. Interestingly, The fitted 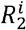 from the sibling control group was relatively higher than that from the MS group (Figure 8C), originally ascribed to an excess iron deposition as a predisposition for MS.^32^ As iron-induced transverse relaxation appears orientation-independent,^20^ it is thus essential to accurately separate 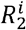 from 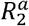 based on a reliable biophysical model for a meaningful evaluation of iron deposition in WM.^70^

### 5.5 Anisotropic *R*_2_ relaxation component

For an anisotropic contribution, its maximum could be 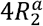as if all the relevant RDCs become consistently orientated along *B*_0_, i.e., *f* (0*°, 0°*) =4. The observed 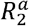 relied on not only the amount of RDCs but also the degree of their restricted molecular motions. For instance, a relatively decreased 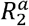 was found in neonatal developing WM (Figure 7A) and compromised WM from people with MS (Figure 8B). Perhaps, an extreme “demyelination” case would be elongated cellular constituents (e.g., microtubules and neurofilaments) in an intra-axonal space, revealing a considerably reduced 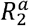 compared to that in an extra-axonal space as shown in Supplementary Figure S4.^7^ On the other hand, a considerably increased 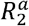 was measured at room temperatures on formalin-fixed WM specimens embedded in an agarose gel at higher fields (≥ 7T).^40, 41^ Moreover, the measured 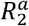 from semisolid CH2 groups in WM was much larger than that from dynamically ordered water close to the phospholipid bilayer surface (Table 1).

It should be clarified that previously published 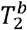 (i.e., ∼10-15 *μs*) from qMT imaging may not be directly linked to the measured CH2 proton resonance absorption linewidth Δ*v* (i.e., ∼3-6 kHz) via the relationship of 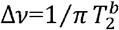, commonly used in an isotropic solution.^57, 71^ Thus, it may not be appropriate to interpret the fitted 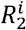 and 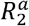 in the same way as those found from their water counterparts. It has been shown before that the same qMT imaging data from bovine WM specimen could be fitted using either Lorentzian 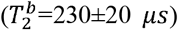 or Super Lorentzian 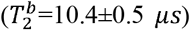 line shape for semisolid protons, albeit with a reduced fitting error for the latter line shape.^72^ If a similar scaling factor (∼25) had been incorporated into the fitted 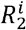 and 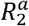 in this work, these values would have been scaled down to the order of kHz as expected. Nonetheless, based on the comparable fitted 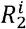 and 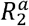, we can conclude that about 80% of 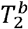 originated from orientation-dependent semisolid CH2 protons in WM, i.e., 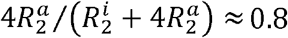.

## 6 CONCLUSIONS

In summary, a cylindrical helix model, encompassing both anisotropic molecular translation and rotation, has been proposed for orientation-dependent proton transverse relaxation in the human brain white matter. This proposed general model can better characterize previously published anisotropic transverse relaxation profiles from various physiological and pathological conditions, thereby providing further insight into myelin microstructural alterations at the molecular level.

## Abbreviations

AR: angular resolution
CPMG: Carr-Purcell-Meiboom-Gill
CNT: control
DTI: diffusion tensor imaging
FA: fractional anisotropy
IE: intra- and extra-cellular
MAE: magic angle effect
MO: mode of anisotropy
MS: multiple sclerosis
PCC: Pearson correlation coefficient
qMT: quantitative magnetization transfer
RDC: residual dipolar coupling
RMSE: root-mean-square error
SBL: siblings
WM: white matter

## ACKNOWLEDGEMENTS

The author is very grateful to Dr. Harald E. Möller (Max Planck Institute for Human Cognitive and Brain Sciences, Leipzig, Germany) for his insightful comments and constructive criticisms, and to Dr. Vladimír Mlynárik (Medical University of Vienna, Vienna, Austria) for his valuable suggestions on the earlier versions of this manuscript. The author would also like to thank Dr. Fuyixue Wang (Massachusetts General Hospital, Charlestown, MA, USA) and Dr. Enedino Hernández-Torres (University of British Columbia, Vancouver, Canada) for sharing Connectome DTI and clinical anisotropic 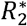 data in public domains.

## DATA AVAILABILITY STATEMENT

Data used in this work are available on request from the author.

## SUPPORTING INFORMATION

**FIGURE S1.**
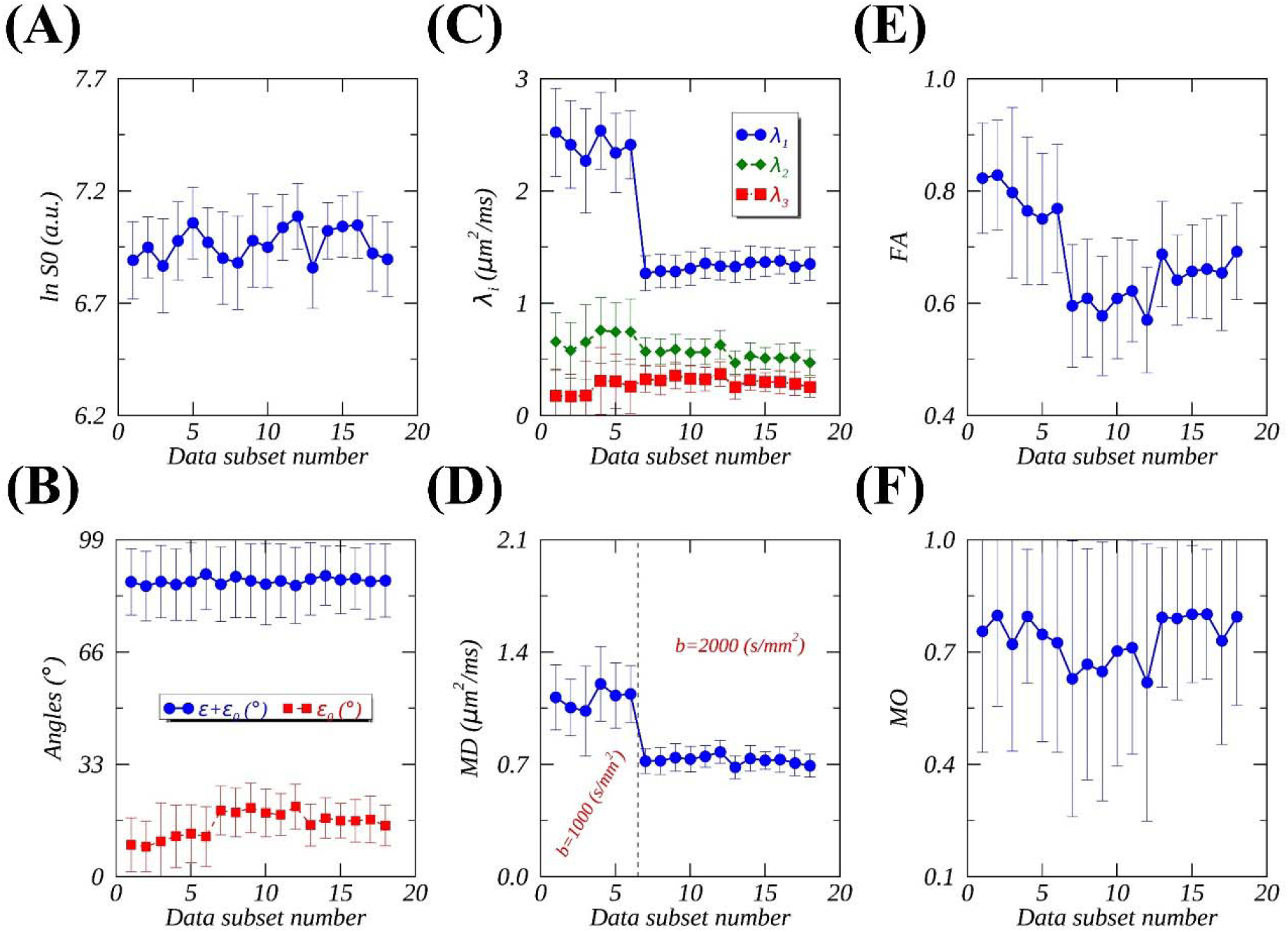
Examples of DTIFIT^48^ output parameters averaged (mean ± SD) within a rectangular ROI from the corpus callosum (as highlighted in Fig. 4D) as well as angle offsets *ε*_0_ (i.e., tan^−1^ *D*_⊥_*/D*_‖_) for 18 diffusion weighting data subsets, acquired using *b*-values (s/mm) of 1000 (n=1-6) and 2000 (n=7-18). Axial or principal (*D*_‖_) and radial (*D*_⊥_) diffusivities were defined as *D*_‖_ *= λ*_1_ and *D*_⊥_ *=* (*λ*_2_+ *λ*_3_)/2, assuming an axially symmetric diffusion tensor.

**FIGURE S2.**
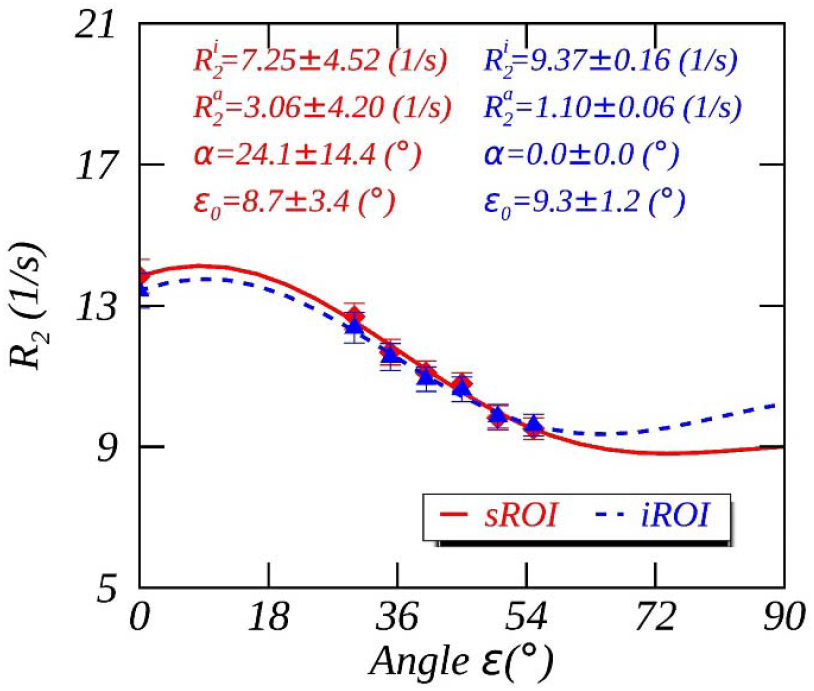
Magic angle effect in peripheral nervous system.^58^ Anisotropic *R* orientation dependences modeled using Fit A for two ROIs: (1) interfascicular space (sROI, red solid line) and (2) interfascicular space and intraneural nerve fascicles (iROI, blue dashed line).

**FIGURE S3.**
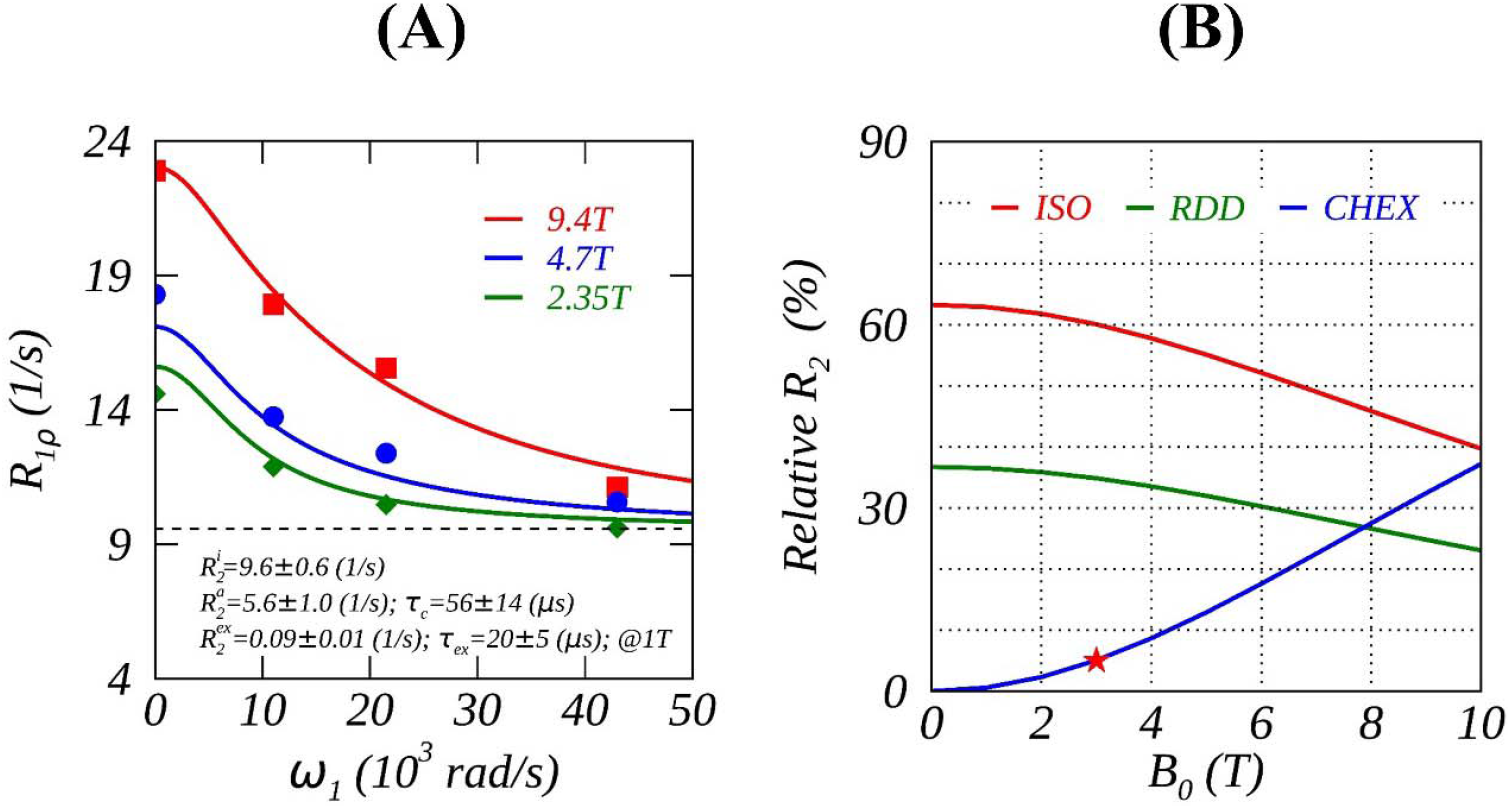
Separating *B*_0_ dependent *R*_1*ρ*_ dispersion^59^ (A) into different contributions (B) on rat brain. *R*_1*ρ*_ comprised a constant (ISO) and two dispersed components: (1) chemical exchange effect (CHEX) and (2) residual dipolar interactions (RDD) as demonstrated previously.^60^ All orientation-independent and *B*_0_-dependent relaxation mechanisms (including susceptibility anisotropy) were lumped into CHEX that is proportional to 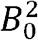. When a spin-lock amplitude *ω*_1_ *=* 0, *R*_1*ρ*_ became *R*_2_ to which the relative CHEX contribution, i.e., 100*CHEX/(ISO+CHEX+RDD), was about 5% at 3T (red star, B).

**FIGURE S4.**
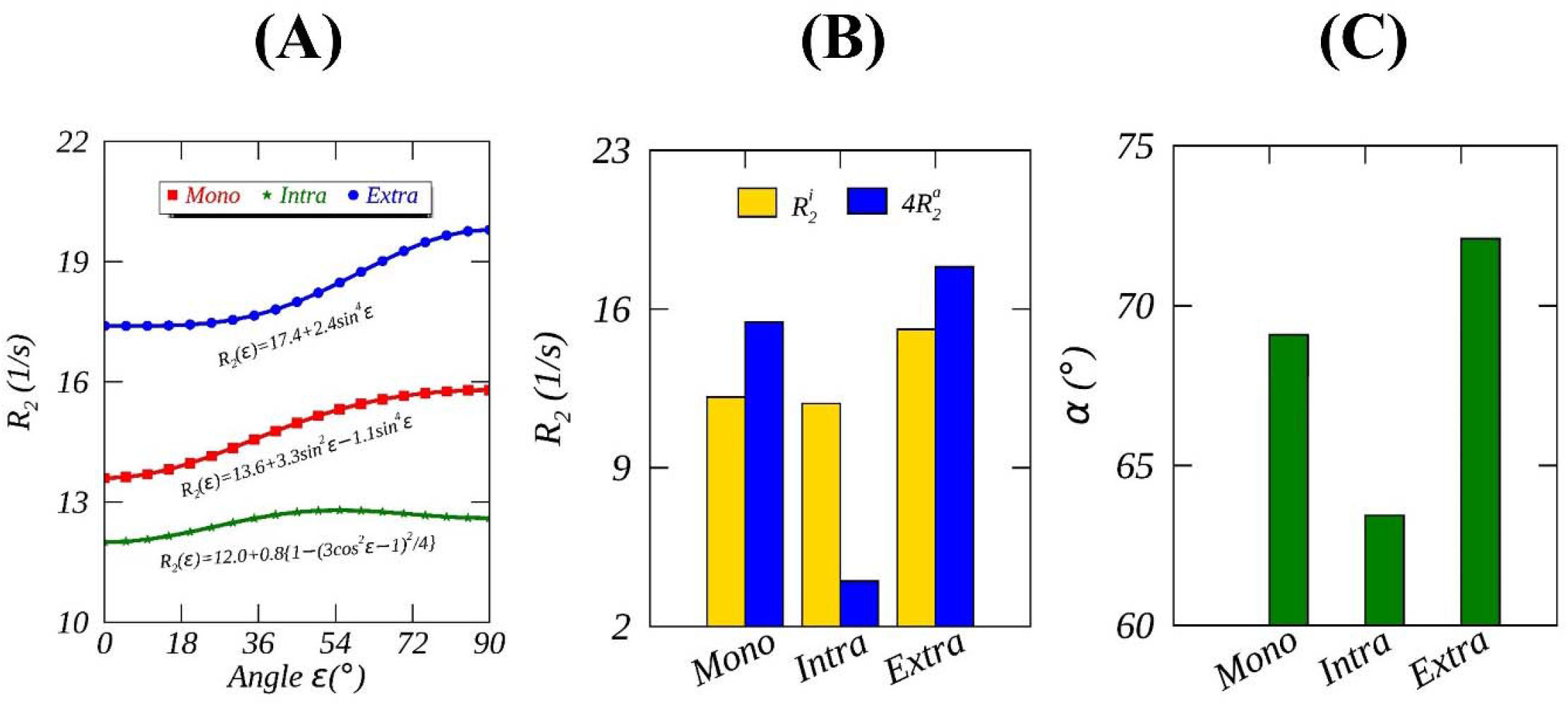
Recast best fitting functions (colored symbols, A) of mono-exponential (red), intra-(green) and extra-axonal (blue) transverse relaxation orientation dependences in WM into 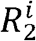 and 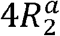 (B), and an open angle *α* (C) using Fit A (colored solid lines, A). The original best fits were obtained from five healthy subjects, using a tiltable RF coil and diffusion-*T*_2_ correlation MRI on a Connectome scanner.^7^

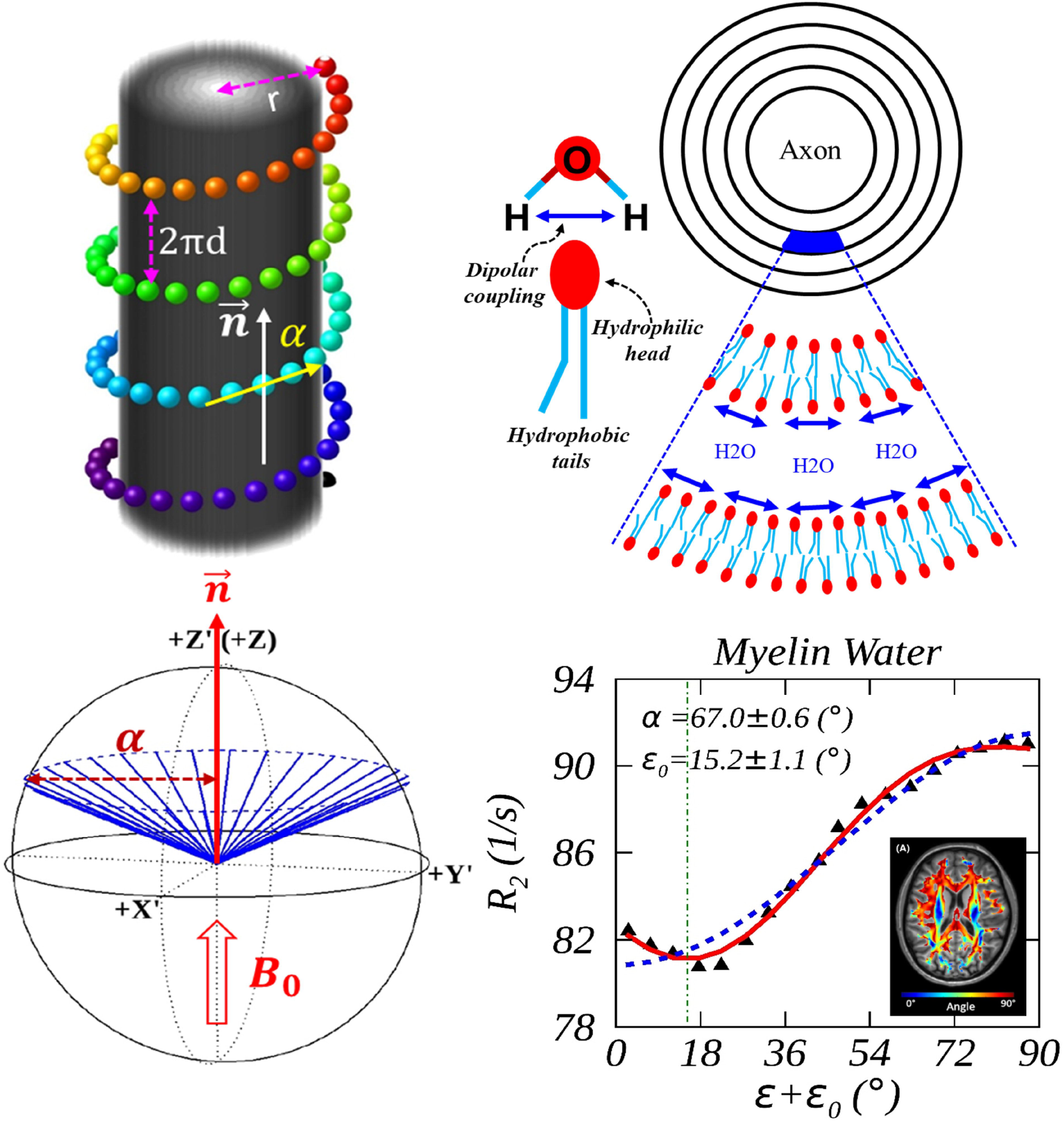

## REFERENCES

1. Chávez FV, Halle B. Molecular basis of water proton relaxation in gels and tissue. Magn Reson Med. 2006;56(1):73–81.

2. Knispel RR, Thompson RT, Pintar MM. Dispersion of proton spin-lattice relaxation in tissues. J Magn Reson. 1974;14(1):44–51.

3. Persson E, Halle B. Cell water dynamics on multiple time scales. Proc Natl Acad Sci USA. 2008;105(17):6266–6271.

4. Schyboll F, Jaekel U, Weber B, Neeb H. The impact of fibre orientation on T 1- relaxation and apparent tissue water content in white matter. Magnetic Resonance Materials in Physics, Biology and Medicine. 2018;31(4):501–510.

5. Schyboll F, Jaekel U, Petruccione F, Neeb H. Origin of orientation-dependent R1 (= 1/T1) relaxation in white matter. Magnet Reson Med. 2020;84(5):2713–2723.

6. Lee J, Nam Y, Choi JY, Kim EY, Oh SH, Kim DH. Mechanisms of T2* anisotropy and gradient echo myelin water imaging. NMR Biomed. 2017;30(4):e3513.

7. Tax CMW, Kleban E, Chamberland M, Baraković M, Rudrapatna U, Jones DK. Measuring compartmental T(2)-orientational dependence in human brain white matter using a tiltable RF coil and diffusion-T(2) correlation MRI. Neuroimage. 2021;236:117967.

8. Basser PJ, Pierpaoli C. Microstructural and physiological features of tissues elucidated by quantitative-diffusion-tensor MRI. J Magn Reson. 2011;213(2):560–570.

9. Beaulieu C. The biological basis of diffusion anisotropy. Diffusion MRI. Elsevier; 2014:155–183.

10. Le Bihan D, Mangin JF, Poupon C, et al. Diffusion tensor imaging: concepts and applications. J Magn Reson Imaging. 2001;13(4):534–546.

11. Dibb R, Liu C. Joint eigenvector estimation from mutually anisotropic tensors improves susceptibility tensor imaging of the brain, kidney, and heart. Magnet Reson Med. 2017;77(6):2331–2346.

12. Bao L, Xiong C, Wei W, Chen Z, van Zijl PC, Li X. Diffusion-regularized susceptibility tensor imaging (DRSTI) of tissue microstructures in the human brain. Medical image analysis. 2021;67:101827.

13. Wisnieff C, Liu T, Spincemaille P, Wang S, Zhou D, Wang Y. Magnetic susceptibility anisotropy: cylindrical symmetry from macroscopically ordered anisotropic molecules and accuracy of MRI measurements using few orientations. Neuroimage. 2013;70:363–376.

14. Wharton S, Bowtell R. Gradient echo based fiber orientation mapping using R2* and frequency difference measurements. Neuroimage. 2013;83:1011–1023.

15. Georgiadis M, Schroeter A, Gao Z, et al. Nanostructure-specific X-ray tomography reveals myelin levels, integrity and axon orientations in mouse and human nervous tissue. Nature communications. 2021;12(1):1–13.

16. Denk C, Torres EH, MacKay A, Rauscher A. The influence of white matter fibre orientation on MR signal phase and decay. NMR Biomed. 2011;24(3):246–252.

17. Weber AM, Zhang Y, Kames C, Rauscher A. Myelin water imaging and R2* mapping in neonates: Investigating R2* dependence on myelin and fibre orientation in whole brain white matter. NMR Biomed. 2020;33(3):e4222.

18. Birkl C, Doucette J, Fan M, Hernandez-Torres E, Rauscher A. Myelin water imaging depends on white matter fiber orientation in the human brain. Magn Reson Med. 2021;85(4):2221–2231.

19. Lee J, van Gelderen P, Kuo L-W, Merkle H, Silva AC, Duyn JH. T2*-based fiber orientation mapping. Neuroimage. 2011;57(1):225–234.

20. Oh S-H, Kim Y-B, Cho Z-H, Lee J. Origin of B0 orientation dependent R2*(= 1/T2*) in white matter. Neuroimage. 2013;73:71–79.

21. Knight MJ, Dillon S, Jarutyte L, Kauppinen RA. Magnetic resonance relaxation anisotropy: Physical principles and uses in microstructure imaging. Biophysical journal. 2017;112(7):1517–1528.

22. Knight MJ, Wood B, Couthard E, Kauppinen R. Anisotropy of spin-echo T2 relaxation by magnetic resonance imaging in the human brain in vivo. Biomedical Spectroscopy and Imaging. 2015;4(3):299–310.

23. Cheng J-X, Pautot S, Weitz DA, Xie XS. Ordering of water molecules between phospholipid bilayers visualized by coherent anti-Stokes Raman scattering microscopy. Proc Natl Acad Sci USA. 2003;100(17):9826–9830.

24. Berendsen HJC. Nuclear magnetic resonance study of collagen hydration. J Chem Phys. 1962;36(12):3297–3305.

25. Pang Y. Characterization of anisotropic T2W signals from human knee femoral cartilage: The magic angle effect on a spherical surface. NMR Biomed. 2021;34(7):e4535.

26. Novikov DS, Kiselev VG, Jespersen SN. On modeling. Magnet Reson Med. 2018;79(6):3172–3193.

27. Veraart J, Novikov DS, Fieremans E. TE dependent Diffusion Imaging (TEdDI) distinguishes between compartmental T2 relaxation times. NeuroImage. 2018;182:360–369.

28. Slator PJ, Palombo M, Miller KL, et al. Combined diffusion-relaxometry microstructure imaging: Current status and future prospects. Magnet Reson Med. 2021;86(6):2987–3011.

29. Bloom M, Burnell EE, MacKay AL, Nichol CP, Valic MI, Weeks G. Fatty acyl chain order in lecithin model membranes determined from proton magnetic resonance. Biochemistry. 1978;17(26):5750–5762.

30. Watnick PI, Dea P, Chan SI. Characterization of the transverse relaxation rates in lipid bilayers. Proc Natl Acad Sci USA. 1990;87(6):2082–2086.

31. Pampel A, Müller DK, Anwander A, Marschner H, Möller HE. Orientation dependence of magnetization transfer parameters in human white matter. Neuroimage. 2015;114:136–146.

32. Hernández-Torres E, Wiggermann V, Hametner S, et al. Orientation dependent MR signal decay differentiates between people with MS, their asymptomatic siblings and unrelated healthy controls. Plos One. 2015;10(10):e0140956.

33. Mathur-De Vre R. The NMR studies of water in biological systems. Progress in Biophysics and Molecular Biology. 1980;35:103–134.

34. Woessner DE. Nuclear magnetic-relaxation and structure in aqueous heterogenous systems. Mol Phys. 1977;34(4):899–920.

35. Lenk R, Bonzon M, Greppin H. Dynamically oriented biological water as studied by NMR. Chemical Physics Letters. 1980;76(1):175–177.

36. Pang Y. An order parameter without magic angle effect (OPTIMA) derived from R1ρ dispersion in ordered tissue. Magn Reson Med. 2020;83(5):1783–1795.

37. Wennerström H. Proton nuclear magnetic resonance lineshapes in lamellar liquid crystals. Chemical Physics Letters. 1973;18(1):41–44.

38. Bydder M, Rahal A, Fullerton GD, Bydder GM. The magic angle effect: a source of artifact, determinant of image contrast, and technique for imaging. J Magn Reson Imaging. 2007;25(2):290–300.

39. Grunder W. MRI assessment of cartilage ultrastructure. NMR Biomed. 2006;19(7):855– 876.

40. Aggarwal M, Kageyama Y, Li X, Van Zijl PC. B0-orientation dependent magnetic susceptibility-induced white matter contrast in the human brainstem at 11.7 T. Magnet Reson Med. 2016;75(6):2455–2463.

41. Pang Y. Anisotropic transverse relaxation in the human brain white matter induced by restricted rotational diffusion. In: Proceedings of the International Society of Magnetic Resonance Medicine, Virtual, 2021:1711.

42. Bender B, Klose U. The in vivo influence of white matter fiber orientation towards B0 on T2* in the human brain. NMR Biomed. 2010;23(9):1071–1076.

43. Doucette J, Wei L, Hernandez-Torres E, et al. Rapid solution of the Bloch-Torrey equation in anisotropic tissue: Application to dynamic susceptibility contrast MRI of cerebral white matter. Neuroimage. 2019;185:198–207.

44. Wharton S, Bowtell R. Fiber orientation-dependent white matter contrast in gradient echo MRI. Proc Natl Acad Sci USA. 2012;109(45):18559–18564.

45. Hédouin R, Metere R, Chan K-S, et al. Decoding the microstructural properties of white matter using realistic models. NeuroImage. 2021;237:118138.

46. Adelnia F, Zu Z, Spear JT, Weng F, Harkins KD, Gore JC. Tissue characterization using R1rho dispersion imaging at low locking fields. Magnetic Resonance Imaging. 2021;84:1–11.

47. Wang F, Dong Z, Tian Q, et al. In vivo human whole-brain Connectom diffusion MRI dataset at 760 μm isotropic resolution. Scientific Data. 2021;8(1):1–12.

48. Smith SM, Jenkinson M, Woolrich MW, et al. Advances in functional and structural MR image analysis and implementation as FSL. Neuroimage. 2004;23:S208–S219.

49. Ennis DB, Kindlmann G. Orthogonal tensor invariants and the analysis of diffusion tensor magnetic resonance images. Magn Reson Med. 2006;55(1):136–146.

50. Pang Y, Palmieri-Smith RM, Malyarenko DI, Swanson SD, Chenevert TL. A unique anisotropic R2 of collagen degeneration (ARCADE) mapping as an efficient alternative to composite relaxation metric (R2 -R1 rho) in human knee cartilage study. Magn Reson Med. 2019;81(6):3763–3774.

51. Papazoglou S, Streubel T, Ashtarayeh M, et al. Biophysically motivated efficient estimation of the spatially isotropic component from a single gradient-recalled echo measurement. Magnet Reson Med. 2019;82(5):1804–1811.

52. Mädler B, Drabycz SA, Kolind SH, Whittall KP, MacKay AL. Is diffusion anisotropy an accurate monitor of myelination?: Correlation of multicomponent T2 relaxation and diffusion tensor anisotropy in human brain. Magnetic resonance imaging. 2008;26(7):874–888.

53. Bevington PR, Robinson DK. Data reduction and error analysis for the physical sciences. 3rd ed. McGraw-Hill; 2003.

54. Markwardt CB. In: Bohlender D, Dowler P, Durand D, eds. Proceedings of Astronomical Data Analysis Software and Systems XVIII, Quebec, Canada, ASP Conference Series, Vol. 411. San Francisco, CA: Astronomical Society of the Pacific; 2009:251–254.

55. Ahearn TS, Staff RT, Redpath TW, Semple SIK. The use of the Levenberg–Marquardt curve-fitting algorithm in pharmacokinetic modelling of DCE-MRI data. Physics in Medicine & Biology. 2005;50(9):N85.

56. Williams L-A, Gelman N, Picot PA, et al. Neonatal brain: regional variability of in vivo MR imaging relaxation rates at 3.0 T—initial experience. Radiology. 2005;235(2):595–603.

57. Seiter C, Chan SI. Molecular motion in lipid bilayers. Nuclear magnetic resonance line width study. Journal of the American Chemical Society. 1973;95(23):7541–7553.

58. Kästel T, Heiland S, Bäumer P, Bartsch A, Bendszus M, Pham M. Magic angle effect: a relevant artifact in MR neurography at 3T? American journal of neuroradiology. 2011;32(5):821–827.

59. Mäkelä HI, De Vita E, Gröhn OH, et al. B0 dependence of the on-resonance longitudinal relaxation time in the rotating frame (T1ρ) in protein phantoms and rat brain in vivo. Magn Reson Med. 2004;51(1):4–8.

60. Pang Y, Palmieri-Smith RM, Maerz T. An efficient R1ρ dispersion imaging method for human knee cartilage using constant magnetization prepared turbo-FLASH. NMR Biomed. 2021;34(6):e4500.

61. Lebel C, Gee M, Camicioli R, Wieler M, Martin W, Beaulieu C. Diffusion tensor imaging of white matter tract evolution over the lifespan. Neuroimage. 2012;60(1):340–352.

62. Bartels L, Doucette J, Birkl C, Zhang Y, Weber AM, Rauscher A. Orientation dependence of T2 in newborn white matter shows dipole-dipole interaction effects. In: Proceedings of the International Society of Magnetic Resonance Medicine, Virtual, 2021:2254.

63. Ohvo-Rekilä H, Ramstedt B, Leppimäki P, Slotte JP. Cholesterol interactions with phospholipids in membranes. Progress in lipid research. 2002;41(1):66–97.

64. Knight MJ, Smith-Collins A, Newell S, Denbow M, Kauppinen RA. Cerebral white matter maturation patterns in preterm infants: an MRI T2 relaxation anisotropy and diffusion tensor imaging study. Journal of Neuroimaging. 2018;28(1):86–94.

65. Bloom M, Sternin E. Transverse nuclear spin relaxation in phospholipid bilayer membranes. Biochemistry. 1987;26(8):2101–2105.

66. Veraart J, Nunes D, Rudrapatna U, et al. Noninvasive quantification of axon radii using diffusion MRI. Elife. 2020;9:e49855.

67. Borthakur A, Wheaton AJ, Gougoutas AJ, et al. In vivo measurement of T1ρ dispersion in the human brain at 1.5 tesla. J Magn Reson Imaging. 2004;19(4):403–409.

68. Momot KI, Pope JM, Wellard RM. Anisotropy of spin relaxation of water protons in cartilage and tendon. NMR Biomed. 2010;23(3):313–24.

69. Mitsumori F, Watanabe H, Takaya N, et al. Toward understanding transverse relaxation in human brain through its field dependence. Magnet Reson Med. 2012;68(3):947–953.

70. Möller HE, Bossoni L, Connor JR, et al. Iron, myelin, and the brain: neuroimaging meets neurobiology. Trends in neurosciences. 2019;42(6):384–401.

71. Gent M, Prestegard J. Nuclear magnetic relaxation and molecular motion in phospholipid bilayer membranes. J Magn Reson. 1977;25(2):243–262.

72. Morrison C, Mark Henkelman R. A model for magnetization transfer in tissues. Magnet Reson Med. 1995;33(4):475–482.

